# Cryo-EM structure of the MgtE Mg^2+^ channel pore domain in Mg^2+^-free conditions reveals cytoplasmic pore opening

**DOI:** 10.1101/2020.08.27.270991

**Authors:** Fei Jin, Minxuan Sun, Takashi Fujii, Yurika Yamada, Jin Wang, Andrés D. Maturana, Miki Wada, Shichen Su, Jinbiao Ma, Hironori Takeda, Tsukasa Kusakizako, Atsuhiro Tomita, Yoshiko Nakada-Nakura, Kehong Liu, Tomoko Uemura, Yayoi Nomura, Norimichi Nomura, Koichi Ito, Osamu Nureki, Keiichi Namba, So Iwata, Ye Yu, Motoyuki Hattori

## Abstract

MgtE is a Mg^2+^ channel conserved in organisms ranging from prokaryotes to eukaryotes, including humans, and plays an important role in Mg^2+^ homeostasis. The previously determined MgtE structures in the Mg^2+^-bound, closed state and structure-based functional analyses of MgtE revealed that the binding of Mg^2+^ ions to the MgtE cytoplasmic domain induces channel inactivation to maintain Mg^2+^ homeostasis. However, due to the lack of a structure of the MgtE channel, including its transmembrane domain in Mg^2+^-free conditions, the pore-opening mechanism of MgtE has remained unclear.

Here, we determined the cryoelectron microscopy (cryo-EM) structure of the MgtE-Fab complex in the absence of Mg^2+^ ions. The Mg^2+^-free MgtE transmembrane domain structure and its comparison with the Mg^2+^-bound, closed-state structure, together with functional analyses, showed the Mg^2+^-dependent pore opening of MgtE on the cytoplasmic side and revealed the kink motions of the TM2 and TM5 helices at the glycine residues, which are important for channel activity. Overall, our work provides structure-based mechanistic insights into the channel gating of MgtE.

## INTRODUCTION

The Mg^2+^ ion is a biologically essential element involved in various physiological processes, including the catalytic action of numerous enzymes, the utilization and synthesis of ATP, and the stabilization of RNA and DNA (Hartwig 2001)(Cowan 2002)(Maguire and Cowan 2002). Therefore, homeostatic imbalance of Mg^2+^ ions causes multiple diseases, such as cardiovascular disease, high blood pressure, osteoporosis, and diabetes (Alexander et al. 2008). Accordingly, Mg^2+^ homeostasis is a requisite for all living organisms, and Mg^2+^ channels and transporters are pivotal in Mg^2+^ homeostasis (Groisman et al. 2013)(Romani 2011)(Moomaw and Maguire 2008). MgtE belongs to a family of Mg^2+^ channels that is ubiquitously distributed in eukaryotes, eubacteria and archaea (Smith et al. 1995)(Townsend et al. 1995)(Goytain and Quamme 2005)(Sahni and Scharenberg 2013)(Schweigel-Röntgen and Kolisek 2014). The bacterial MgtE is a Mg^2+^-selective ion channel involved in Mg^2+^ homeostasis (Dann et al. 2007)(Hattori et al. 2009) and is also implicated in bacterial survival after antibiotic exposure (Lee et al. 2019). The human orthologs of MgtE, the SLC41 proteins (SLC41A1, SLC41A2 and SLC41A3), also transport Mg^2+^ ions (Goytain and Quamme 2005)(Mandt et al. 2011)(Sahni et al. 2007)(de Baaij et al. 2016), and mutations in these proteins are associated with diabetes (Chan et al. 2015), Parkinson’s disease (Kolisek et al. 2013), and nephronophthisis (Hurd et al. 2013).

The previously determined crystal structures of MgtE from *Thermus thermophilus* revealed the homodimeric architecture of MgtE, consisting of transmembrane (TM) and cytoplasmic domains connected via a long soluble helix, named the “plug” (Hattori et al. 2007). A high-resolution structure of the MgtE TM domain revealed the ion selectivity filter of MgtE with the fully hydrated Mg^2+^ ion in the ion-conducting pore (Takeda et al. 2014). The cytoplasmic domain of MgtE is composed of two subdomains, namely, the N and Cystathionine-beta-synthase (CBS) domains. In the full-length MgtE structure in the presence of Mg^2+^ ions, the binding of multiple Mg^2+^ ions at the subunit interfaces of the cytoplasmic domain appears to stabilize the closed conformation of MgtE by fixing the orientation of the plug helices to close the ion-conducting pore (**Fig. 1A**). The crystal structure of the Mg^2+^-free cytoplasmic domain exhibits a more relaxed conformation, where the N domain is dissociated from the CBS domain and where each subunit of the CBS domain and plug helix dimer are also separated from each other, which may unlock the closed conformation of the channel (**Fig. 1B and 1C**). Consistent with this, electrophysiological recording of MgtE showed that a high concentration of cellular Mg^2+^ inhibits the channel gating of MgtE, whereas MgtE is in an equilibrium between the open and closed states of the channel under a low concentration of cellular Mg^2+^ (Hattori et al. 2009). However, due to the lack of a MgtE TM domain structure in Mg^2+^-free conditions, the Mg^2+^-dependent structural changes in MgtE, particularly those in the transmembrane domain, remain unknown, and thus, the channel-opening mechanisms remain unclear.

**Figure 1.**
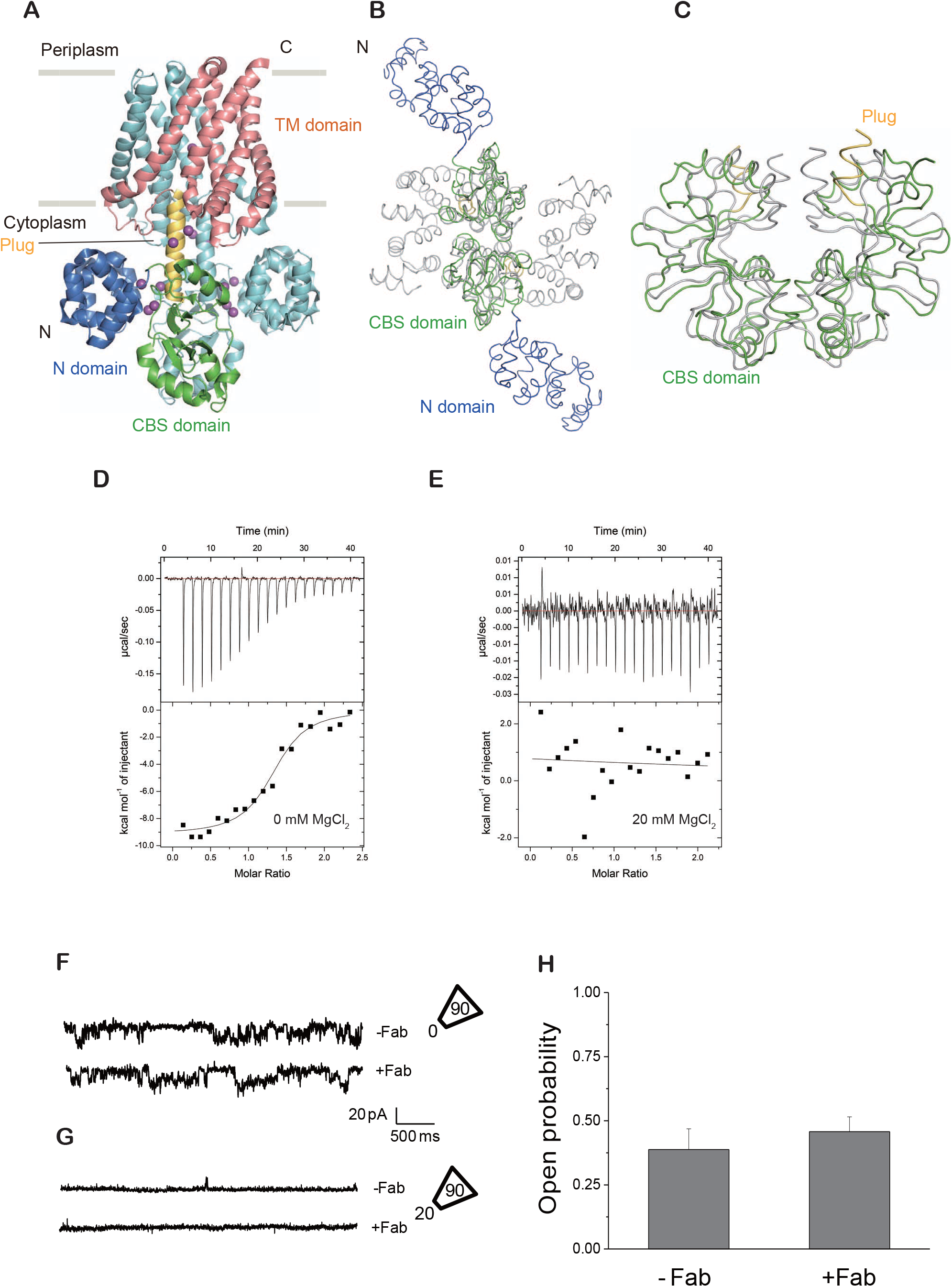
Functional characterization of Fab705. (A) MgtE dimer in the presence of Mg^2+^ (PDB ID:2ZY9), viewed parallel to the membrane. The N and CBS domains, plug helix and TM domain in chain A are colored blue, green, yellow and red, respectively. The other subunit (chain B) is colored cyan. Mg^2+^ ions are depicted as purple spheres. **(B, C)** Structural comparison of the Mg^2+^-bound and Mg^2+^-free MgtE cytoplasmic domains superimposed on the CBS domains and viewed from the cytoplasmic side (**B**) and parallel to the membrane (**C**). **(D, E)** ITC analysis of MgtE with Fab705 in the presence of 0 mM MgCl_2_ (D) and 20 mM MgCl_2_ (E). **(F, G)** Single-channel recording of MgtE in the inside-out configuration from *E. coli* giant spheroplasts. Representative current traces from triple-knockout *E. coli* spheroplasts expressing MgtE in the presence (lower) or absence (upper) of 2 μM Fab705. (**H**). The open probabilities for MgtE with (+) and without (-) 2 μM Fab705 (n=6 for each condition). The error bars represent the SEM.

Here, we determined the cryoelectron microscopy (cryo-EM) structure of the MgtE-Fab complex in the absence of Mg^2+^ ions. The Mg^2+^-free MgtE structure revealed the cytoplasmic pore opening motions of the MgtE TM domain. Structure-based functional analyses further clarified these structural changes on the cytoplasmic side, providing mechanistic insights into the channel gating of MgtE.

## RESULTS

### Generation of the Fab antibody for MgtE

Many years of efforts to crystallize MgtE in Mg^2+^-free conditions yielded only poorly diffracting crystals. Thus, we decided to employ single-particle cryo-EM to determine the structure of MgtE in Mg^2+^-free conditions. However, it was still a challenge to obtain a high-resolution structure of full-length MgtE by cryo-EM due to its low molecular weight. Therefore, we attempted to generate Fab antibodies for MgtE to enable determination of the cryo-EM structure of the MgtE-Fab complex by increasing its total molecular weight.

After mouse immunization and generation of multiple monoclonal antibodies for MgtE, fluorescence-detection size-exclusion chromatography (FSEC) analysis showed that one of the Fab fragments, Fab705, bound to full-length MgtE at 0 mM Mg^2+^, as indicated by the apparent peak shift (**Fig. S1A**), but the binding did not occur at a high concentration of Mg^2+^ ions (20 mM) (**Fig. S1B**). Consistent with this, isothermal titration calorimetry (ITC) analysis showed that Fab705 bound to full-length MgtE at 0 mM MgCl_2_ with a K_*d*_ of 252.6±71.1 nM, whereas no interaction was detected between Fab705 and MgtE under a high concentration of Mg^2+^ ions (20 mM) (**Fig. 1D and 1E) (Table S1**). We then performed patch clamp analysis of full-length MgtE using *Escherichia coli* giant spheroplasts at 0 mM MgCl_2_ in the presence and absence of Fab705 and observed little change in the open probabilities upon addition of Fab705 into the bath solution (**Fig. 1F and 1H)**. These results showed that Fab705 binds to Mg^2+^-free MgtE but not to MgtE at high Mg^2+^ concentrations and does not either positively or negatively modulate channel opening at 0 mM Mg^2+^. These functional properties of Fab705 would be desirable for determination of the MgtE structure under Mg^2+^-free conditions because it does not affect the channel gating under Mg^2+^-free conditions, in particular in an inhibitory manner. Thus, the utilization of Fab705 would enable us to capture the gating motions of MgtE by cryo-EM.

### Cryo-EM of the MgtE-Fab complex in Mg^2+^-free conditions

We performed cryo-EM single-particle analysis to determine the structure of the full-length MgtE-Fab705 complex reconstituted in PMAL-C8 amphipol in the presence of 1 mM EDTA and the absence of divalent cations (**Fig. 2, S2 and S3**). A 3D reconstruction of the cryo-EM images was performed to reach an overall resolution of 3.7 Å, with C2 symmetry imposed **(Fig. 2C**), and the cryo-EM density of the TM domain was clear enough to build an atomic model (**Fig. 2D-F and S4**).

**Figure 2.**
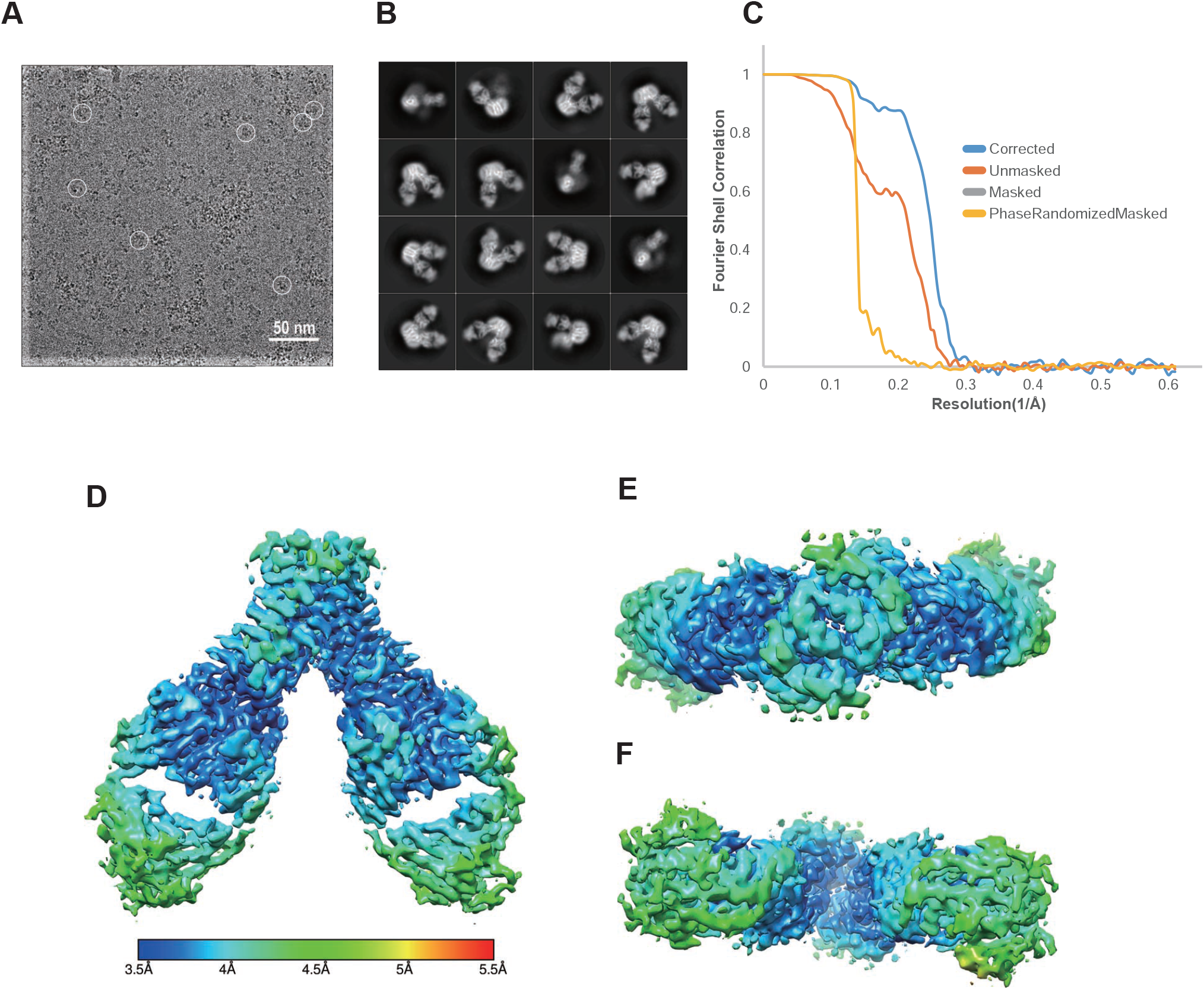
Cryo-EM of the MgtE-Fab complex. **(A)** Cryo-EM image at −1.8 μm defocused with MgtE-Fab complex particles. (**B**) Selected 2D class averages used to produce the 3D reconstruction. (**C**) Gold standard FSC for estimating resolution. (**D-F**) Side and top views of the final map colored according to local resolution, calculated using RELION.

Notably, we did not detect the cryo-EM density of the cytoplasmic domain of MgtE. The SDS-PAGE analysis showed the presence of the cytoplasmic domain of MgtE in the purified sample for Cryo-EM (**Fig. S2**), and thus, the absence of the MgtE cytoplasmic domain in the cryo-EM density was probably because of the high structural flexibility of the Mg^2+^-free cytoplasmic domain (Hattori et al. 2007)(Ishitani et al. 2008). The highly flexible nature of the Mg^2+^-free MgtE cytoplasmic domain in our cryo-EM structure is consistent with previous studies conducted on MgtE by high-speed atomic force microscopy (HS-AFM) (Haruyama et al. 2019) and molecular dynamics (MD) simulations (Ishitani et al. 2008). In particular, the HS-AFM results directly visualized the continuous shaking motions of the MgtE cytoplasmic domains under Mg^2+^-free conditions. The flexibility of the MgtE cytoplasmic domain in Mg^2+^-free conditions is necessary for unlocking the closed conformation of the MgtE transmembrane domain for channel opening (Hattori et al. 2009).

Nevertheless, the details of the structural changes in the TM domain following the release of Mg^2+^ ions from the cytoplasmic domain were previously unclear, and we successfully visualized the conformation of the TM domain under Mg^2+^-free conditions for the first time, to the best of our knowledge. MgtE forms a symmetric dimer, and each subunit is composed of five TM helices (**Fig. 3**). The Fab705 molecules are bound to the cytoplasmic side of the MgtE TM domain, and this binding is mediated mainly by electrostatic interactions and hydrogen bonds, primarily involving the residues in the TM2-TM3 linker the TM4-TM5 linker (**Fig. 3A and S5A**). Among these residues involved in Fab binding, Arg345 interacts with Glu217 in the CBS domain, and Asp418 binds to Mg^2+^ ions to bridge the TM domain and plug helices (**Fig. S5B**).

**Figure 3.**
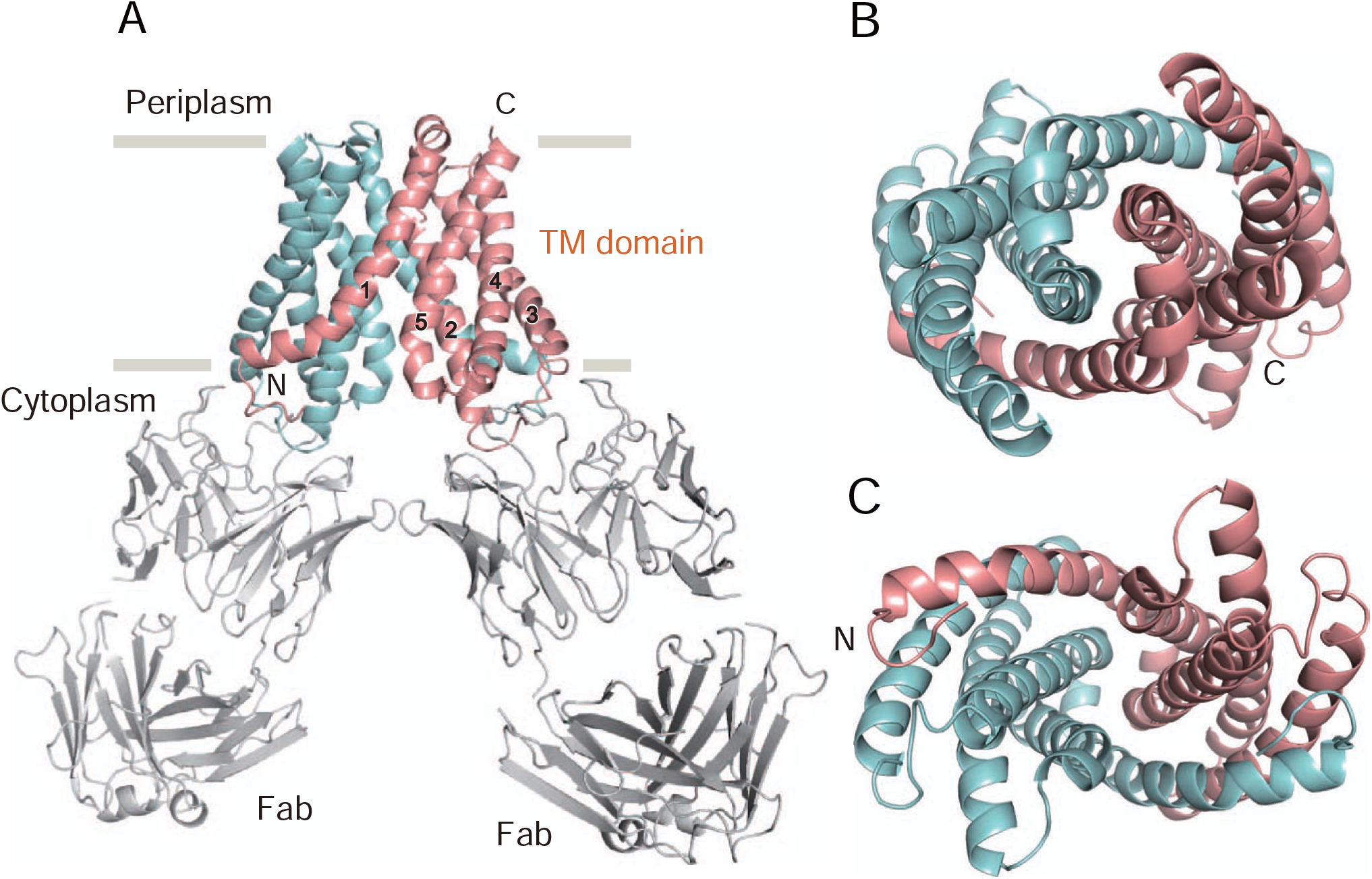
Overall structure. **(A)** Cartoon representation of the MgtE-Fab complex dimer under Mg^2+^-free conditions, viewed parallel to the membrane. The MgtE transmembrane domain dimer is indicated in red (chain A) and cyan (chain B). The Fab fragments are colored in gray. **(B, C)** The MgtE dimer, viewed from the periplasm (B) and from the cytoplasm (C).

The superposition of our cryo-EM structure and the MgtE cytoplasmic domain structure in Mg^2+^-free conditions (PDB ID: 2YVZ) onto the crystal structure of full-length MgtE in the presence of Mg^2+^ (PDB ID: 2ZY9) (**Fig. S6**) reveals the spatial collision between Fabs and the Mg^2+^-bound cytoplasmic domain, mainly at the N domain (**Fig. S6B**), whereas the N domain is dissociated from the CBS domain in the absence of Mg^2+^ (**Fig. S6B**), explaining why we did not observe binding between MgtE and Fab705 at a high concentration of Mg^2+^ ions.

Notably, in the FSEC analysis, the addition of 20 mM Mg^2+^ into the sample of the MgtE-Fab complex sample, which was preformed before the addition of Mg^2+^, disrupted complex formation, where we did not see a Fab-dependent FSEC peak shift (**Fig. S1C**). Consistent with this, in the patch clamp recording, when we tested a high concentration of MgCl_2_ (20 mM MgCl_2_) in the bath solution after the addition of Fab705, we did not observe channel opening, as similarly observed with a high concentration of MgCl_2_ in the absence of Fab705 (**Fig. 1G**). These results indicate that after Fab binding, a high concentration of magnesium ions can still close the MgtE channel by disrupting the MgtE-Fab complex.

We then compared our structure with the previously determined Mg^2+^-bound, closed-state MgtE structure and performed further functional analysis to gain structural insight into MgtE channel opening.

### Ion-conducting pore

The MgtE structure possesses an ion-conducting pore at the center of the dimer (**Fig. 4**). The MgtE structure in Mg^2+^-free conditions showed a wider pore on the cytoplasmic side than that in the previously determined structure in the presence of Mg^2+^ ions (**Fig. 4C**). Notably, at the cytoplasmic end of the pore, the side chains of Asn424 of both subunits face each other in the Mg^2+^-bound closed structure (**Fig. 4B**), whereas in our new Mg^2+^-free structure, the side chains of Asn424 are turned away from each other (**Fig. 4A)** to form the wider cytoplasmic pore, suggesting that this region might form the cytoplasmic gate.

**Figure 4.**
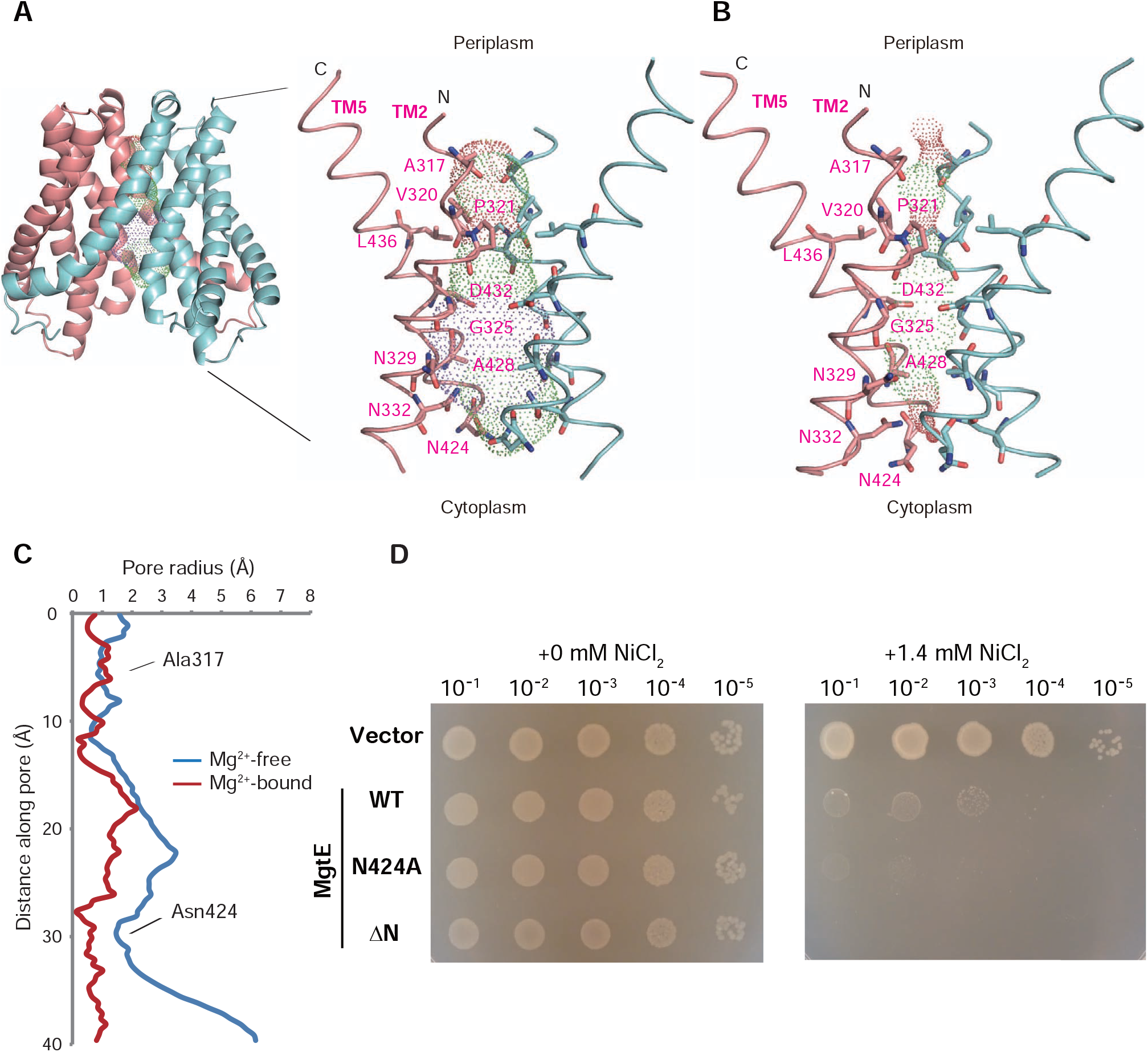
Ion-conducting pore. (**A, B**) The solvent-accessible surface of the MgtE pore under Mg^2+^-free conditions (A) and in the presence of Mg^2+^ (B) with the pore-lining residues. A cartoon representation of the MgtE TM domain dimer is also shown. (**C**) Plots of the pore radius of MgtE structures in Mg^2+^-free and Mg^2+^-bound states. (**D**) Ni^2+^ sensitivity assay of the wild-type *E. coli* strain. Serially diluted cultures (10^−1^, 10^−2^, 10^−3^, 10^−4^, 10^−5^) of magnesium-prototrophic wild-type *E. coli* (W3110 DE3) transformants harboring expression plasmids (empty vector, wild-type MgtE and the N424A and ΔN mutants) were spotted onto the assay LB plates supplemented with 10 μM IPTG (moderate T7 promoter induction): (left) no additional NiCl_2_, (right) + 1.4 mM NiCl_2._

To test this hypothesis, we generated a MgtE mutant possessing an alanine substitution at Asn424, which may disrupt the cytoplasmic gate. We conducted a Ni^2+^ sensitivity assay of *E. coli* with the N424A mutant of MgtE; Mg^2+^ channels, including MgtE, typically transport Ni^2+^ because the coordination chemistry of these ions is highly similar to that of Mg^2+^ (Moomaw and Maguire 2008). Increased sensitivity to Ni^2+^ indicates upregulation of Ni^2+^ uptake, as previously reported with the ΔN mutant of MgtE (Hattori et al. 2009).

*E. coli* cells expressing the N424A mutant of MgtE exhibited increased Ni^2+^ sensitivity (**Fig. 4D**) compared to cells expressing wild-type MgtE, suggesting that disruption of the cytoplasmic gate led to increased Ni^2+^ uptake.

In contrast to the opening of the cytoplasmic side of the pore, the pore remains closed around the Ala317 residues on the periplasmic side, where the pore radius is ∼1 Å (**Fig. 4C**) in both MgtE structures in the presence and absence of Mg^2+^ ions, indicating that the current structure would remain in a nonconductive conformation on the periplasmic side.

### Cytoplasmic pore-opening motions

Consistent with the wider opening of the ion-conducting pore on the cytoplasmic side, the TM helices, including the pore-forming TM2 and TM5 helices, are located in exterior positions with respect to the center of the ion-conducting pore under Mg^2+^-free conditions (**Fig. 5A and 5B) (Video S1**). For instance, compared to the structure of the Mg^2+^-bound state, Thr344 at TM2 and Ala420 at TM5 are rotated by ∼8° and ∼14°, respectively, clockwise from the central axis along the pore and have moved away from the central axis by ∼3 Å and ∼1 Å, respectively. Furthermore, the TM1 helices that are directly connected to the plug helices have also moved away from the center, as they interact with both the TM2 and TM5 helices (**Fig. 5B**).

**Figure 5.**
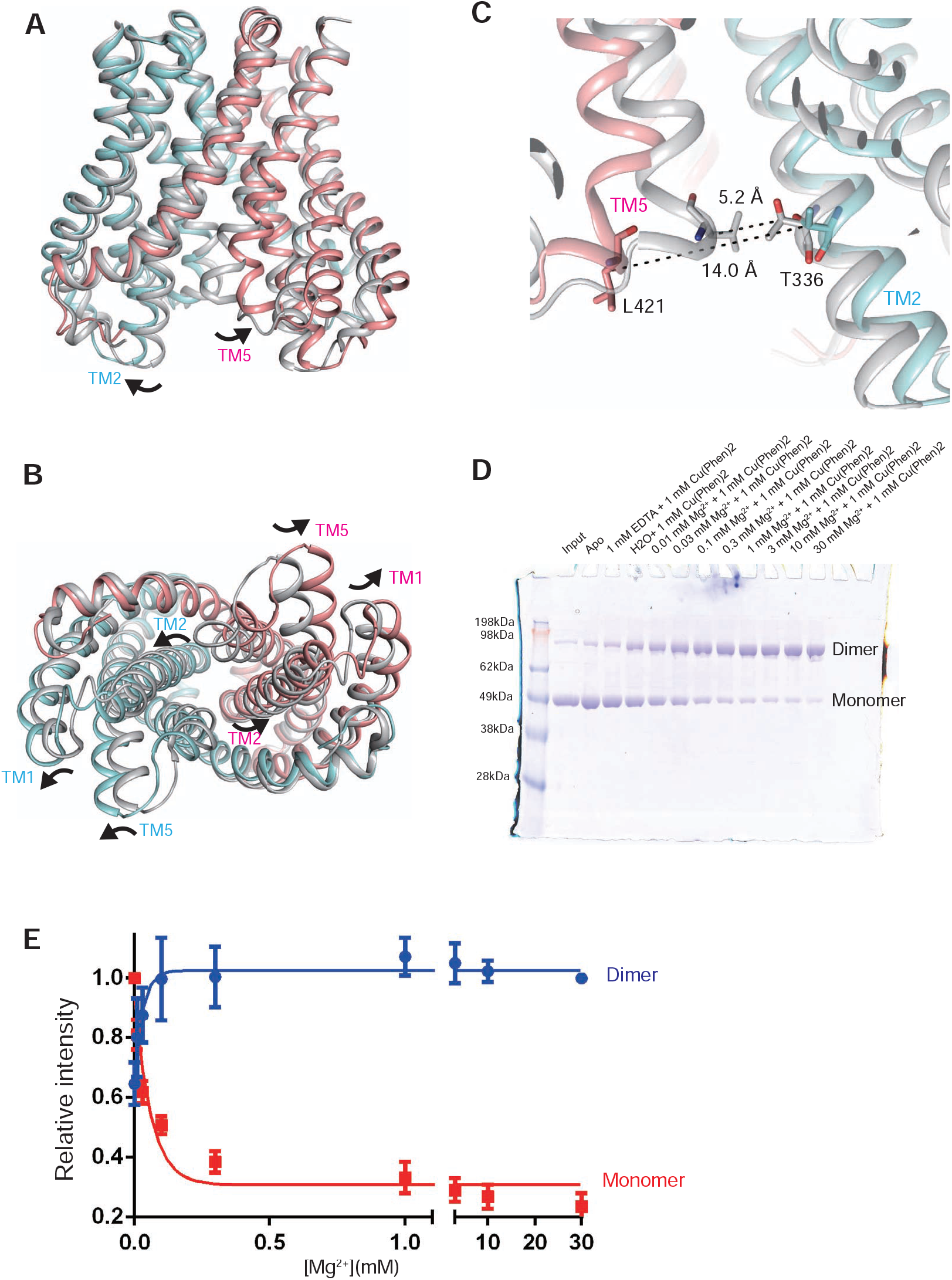
Cytoplasmic pore-opening motions. **(A, B)** Mg^2+^-free MgtE is superposed onto Mg^2+^-bound MgtE in the closed state using the Cα positions of the TM domain dimer, viewed parallel to the membrane (A) and from the cytoplasm (B). Mg^2+^-free MgtE is colored red (chain A) and cyan (chain B). Mg^2+^-bound MgtE is colored gray. The black arrows indicate the structural changes between the Mg^2+^-bound and Mg^2+^-free structures. (**C**) A close-up view of the MgtE dimer interface on the cytoplasmic side. Dotted black lines indicate the Cβ distances between Thr336 and Leu421. (**D**) Chemical cross-linking experiments with the MgtE double cysteine mutant T336C/L421C. (E) Densitometric quantification and sigmoidal curve fitting of SDS-PAGE band intensities from (D). Experiments were repeated six times.

To verify the Mg^2+^-dependent structural changes in the TM helices on the cytoplasmic side, we generated MgtE mutants possessing cysteine substitutions at Leu421 and Thr336, where the Cα distances between Leu421 (A) and Thr336 (B) in the presence and absence of Mg^2+^ ions were 5.2 and 14.0 Å, respectively (**Fig. 5C**). Using these mutants with Cu^2+^ phenanthroline as a catalyst, we performed cross-linking experiments (**Fig. 5D and 5E**). Wild-type MgtE does not contain any endogenous cysteine residues, and the addition of MgCl_2_ to MgtE would shift the equilibrium from the Mg^2+^-free form to the Mg^2+^-bound form, according to the previously reported limited proteolysis assay of MgtE (Hattori et al. 2009). The double cysteine mutant T336C/L421C exhibited a strong band corresponding to the MgtE dimer in the presence of Mg^2+^ at a concentration gradient (0.01-10 mM) and Cu^2+^ phenanthroline (**Fig. 5D and 5E**), consistent with the Mg^2+^-dependent structural changes observed in our cryo-EM structure.

To further support the current conformation of the MgtE TM domain structure, we performed MD simulations starting from the Mg^2+^-free MgtE TM domain structure embedded in the lipid bilayer. The results of the 2-μs simulation showed that the structures of the MgtE TM domain are mostly stable in Mg^2+^-free conditions (**Fig. S7**). The cytoplasmic pore remained open, with Cα distances between Leu421 (A) and Thr336 (B) of 10-15 Å, whereas the periplasmic gate region site remained closed, with stable Cα distances between Ala317 (A) and Ala317 (B) of ∼6 Å during the MD simulations. The results further support the cytoplasmic pore opening of MgtE under Mg^2+^-free conditions, as we observed in the cryo-EM structure. At the same time, we also do not exclude the possibility that we might see some structural changes if we run the MD simulation for a much longer duration.

### Kink motions of the TM helices for channel gating

Detailed observation and comparison of the MgtE structures further revealed the kink motions of TM2 at Gly325 and Gly328 by ∼10° and of TM5 at Gly435 by ∼10° (**Fig. 6A and 6B**). Since the TM2 and TM5 helices form the ion-conducting pore, these kink motions seemingly enable the TM2 and TM5 helices to expand the pore on the cytoplasmic side.

**Figure 6.**
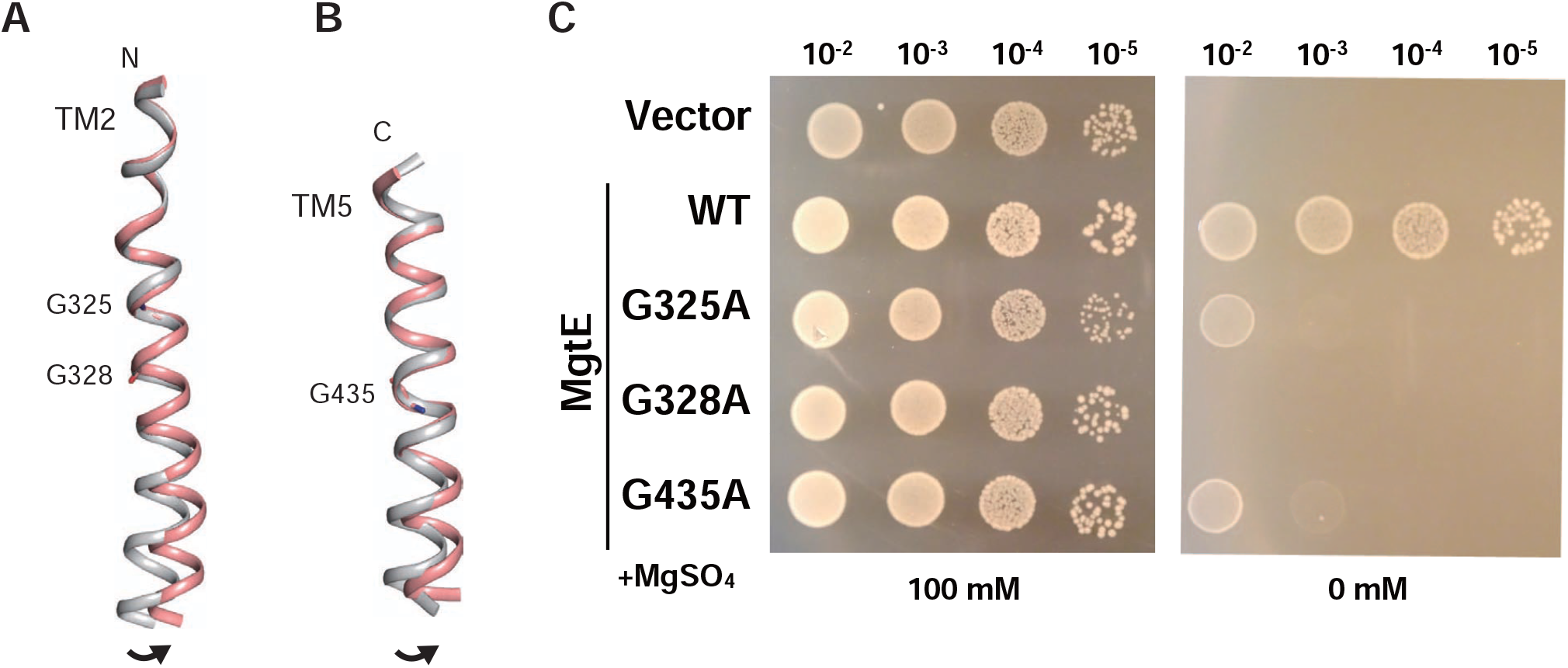
Kink motions of the TM helices. (**A, B**) Structural changes in the TM2 (A) and TM5 (B) helices. The TM helices of Mg^2+^-free MgtE (red) are superimposed on those of Mg^2+^-bound MgtE (gray) using the Cα positions of residues 313–325 for TM2 and 435–447 for TM5. Gly325, Gly328 and Gly435 in Mg^2+^-free MgtE are depicted as stick representations. (**C**) Mg^2+^‐auxotrophic growth complementation assay of MgtE and its mutants on -Mg^2+^ plates. Serially diluted cultures (10^−2^, 10^−3^, 10^−4^, 10^−5^) of Mg^2+^-auxotrophic *E. coli* transformants harboring expression plasmids (vector, wild-type MgtE and the G325A, G328A, and G435A mutants) were spotted onto the assay LB plates.

To examine the functional importance of these glycine residues, we generated a series of alanine-substituted mutants of MgtE (G325A, G328A and G435A) and performed a growth complementation assay using a Mg^2+^-auxotrophic *E. coli* strain lacking the major genes (Δ*mgtA* Δ*corA* Δ*yhiD*) encoding Mg^2+^ transporters that survived only when the medium was supplemented with a sufficiently high concentration of Mg^2+^ (Hattori et al. 2009). The exogenously expressed MgtE could reconstitute the Mg^2+^ transport activity of this strain, and the strain transformed with the wild-type MgtE gene could survive in Mg^2+^-free LB medium (Hattori et al. 2009). While the expression of wild-type MgtE rescued Mg^2+^-auxotrophic growth, the mutations at these glycine residues almost abolished Mg^2+^ transport activity (**Fig. 6C**). In particular, mutation of the strictly conserved Gly328 completely eliminated the growth complementation activity (**Fig. 6C**). These results indicated that these glycine residues at the kinks of the TM2 and TM5 helices are important for the channel activity of MgtE.

## DISCUSSION

In this work, we determined the cryo-EM structure of MgtE in Mg^2+^-free conditions. The utilization of antibodies has been successfully applied to the single-particle analysis of membrane proteins (Lü et al. 2017)(Park et al. 2017)(Manolaridis et al. 2018)(Butterwick et al. 2018)(Coleman et al. 2019). Notably, in our MgtE-Fab complex structure, the ordered region of MgtE (MgtE TM domain dimer) has a molecular weight of only 39 kDa, further demonstrating antibody-based techniques to be powerful tools for determination of membrane protein structures by cryo-EM. To the best of our knowledge, the 39-kDa MgtE TM domain dimer in our study is the smallest membrane protein resolved using single-particle analysis at near-atomic resolution.

Based on this structure, we further conducted structure-based functional analysis, and our new findings together with our previous research on MgtE provided structural insights into the gating mechanism of MgtE, as discussed below.

First, under high Mg^2+^ concentrations, the binding of intracellular Mg^2+^ to the cytoplasmic domain of MgtE, including the plug helices, stabilizes the tightly packed, closed conformation of the protein, leading to pore closure in the TM domain through the extensive interactions between the plug helices and the TM domain, mainly via helices TM2 and TM5 on the cytoplasmic side (Hattori et al. 2007) (**Fig. 1A and 7A**).

**Figure 7.**
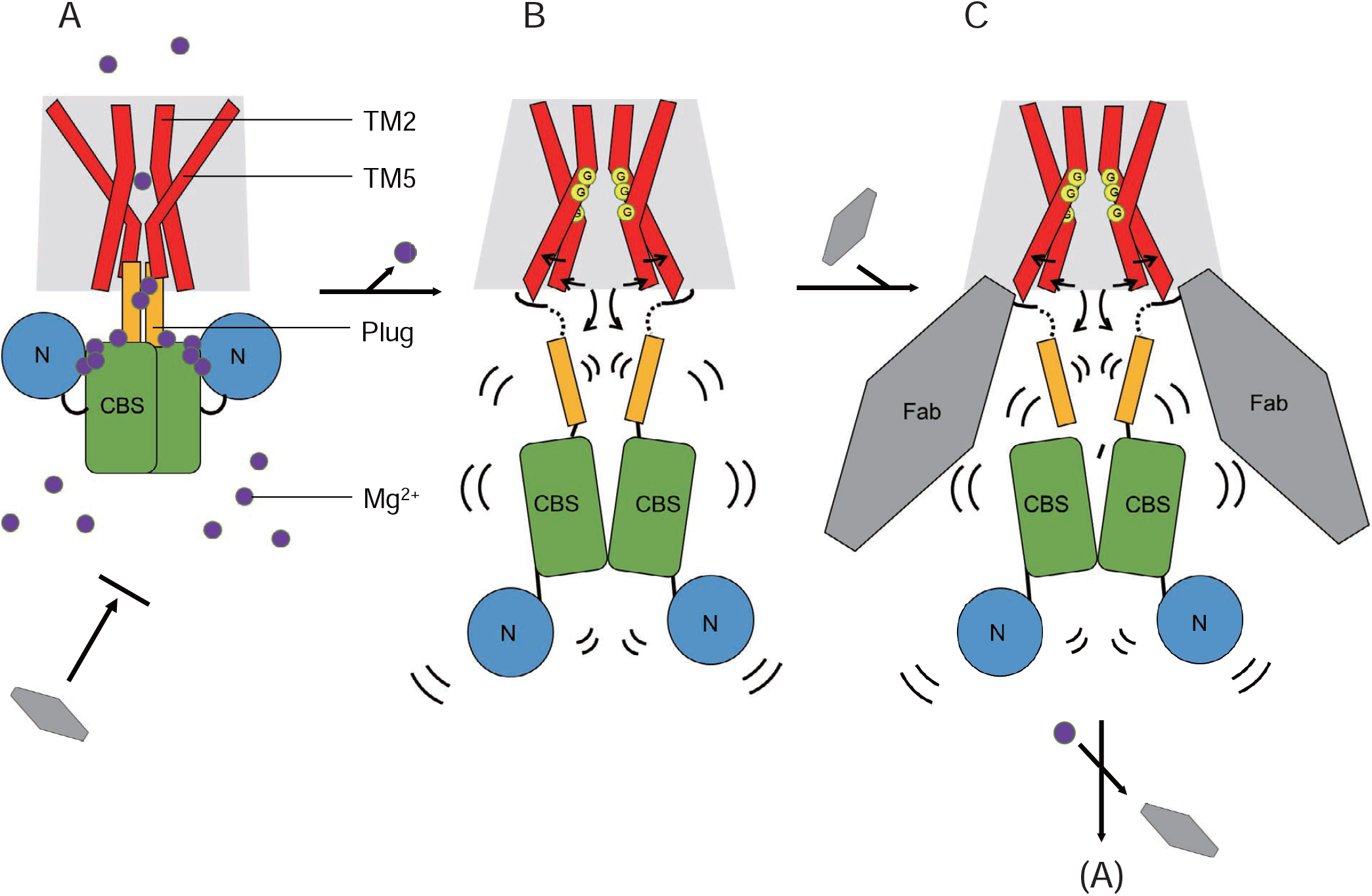
Proposed mechanism. A cartoon of the proposed gating motions in the presence of Mg^2+^ (**A**) and under Mg^2+^-free conditions (**B**). MgtE transmembrane domain, plug helix, CBS domain and N domain are depicted in red, yellow, green and blue, respectively. The purple circles represent magnesium ions. The yellow circles containing G emphasize the kink motions at the glycine residues (Gly325, Gly328 and Gly435). The black arrows indicate the structural changes between the Mg^2+^-bound state and the Mg^2+^-free structures. Curved black lines indicate the flexibility of the cytoplasmic domain under Mg^2+^-free conditions. The binding modes of Fab705 are also presented **(A) (C)**. Briefly, Fab705 cannot bind to MgtE at high Mg^2+^ concentrations (**A**). At low Mg^2+^ concentrations, Fab705 can bind to MgtE, but does not either positively or negatively modulate channel opening, and a high concentration of Mg^2+^ ions can still close the MgtE channel by disrupting the MgtE-Fab complex (**C**).

In contrast, under Mg^2+^-free conditions, the MgtE cytoplasmic domain, including the plug helices, was totally disordered in our cryo-EM map (**Fig. 2**), indicating the highly flexible nature of the MgtE cytoplasmic domain under Mg^2+^-free conditions (**Fig. 7B**). In such disordered conditions, the plug helices of MgtE cannot tightly interact with the TM domain to lock the channel in the closed state any more, unlike the structure in the Mg^2+^-bound state (**Fig. 1A and 7A**). In other words, under Mg^2+^-free conditions, the high flexibility of the cytoplasmic domain leads to loss of the interactions between the plug helices and TM domains, unlocking the TM domain conformation from the closed state. Highly flexible domain motions in the MgtE cytoplasmic domain were also observed by high-speed atomic force microscopy (Haruyama et al. 2019), in the crystal structure of the Mg^2+^-free cytoplasmic domain structure (Hattori et al. 2007), and via MD simulations of MgtE (Ishitani et al. 2008). In particular, in a previous report on HS-AFM (Haruyama et al. 2019), Haruyama and his colleagues observed a jagged topography of MgtE in the absence of Mg^2+^ under HS-AFM from the end-up orientation (cytoplasmic view), indicating that the cytoplasmic domains are constantly fluctuating. Furthermore, they observed that in the side-on orientation (side view), the cytoplasmic regions of MgtE were highly flexible and ambiguous in the absence of Mg^2+^, and successive images demonstrated the association and disassembly of the N and CBS domains in solution. Such disassembly of the N and CBS domains is consistent with the previous structure of the MgtE cytoplasmic domain under Mg^2+^-free conditions (**Fig. 1B and 7B**).

Following the unlocking of the TM domain due to the high flexibility of the Mg^2+^-free cytoplasmic domain, we observed the structural differences of the TM domains between our cryo-EM structure and the previously determined Mg^2+^-bound, inactivated structure (**Fig. 5**). In our cryo-EM structure, the ion-conducting pore is wide open on the cytoplasmic side but still closed on the periplasmic side (**Fig. 4**), indicating that the pore is still nonconductive. According to previous patch clamp recording results, MgtE is in equilibrium between the open and closed states under low-Mg^2+^ conditions rather than in a constitutively open state (Hattori et al. 2009). Therefore, our cryo-EM structure of MgtE may represent a preopening state of MgtE under low-Mg^2+^ conditions, where the pore is partially opened on the cytoplasmic side due to loss of the tight interactions with the plug helices (**Fig. 7B**).

In our structure, the structural changes in the TM domain occur mainly at the TM2 and TM5 helices (**Fig. 5 and 7B**), and the glycine residues in the TM2 and TM5 helices enable the kink motions of the TM2 and TM5 helices for cytoplasmic pore opening (**Fig. 6 and 7B)**. We verified these structural changes by biochemical cross-linking experiments and genetic assays **(Fig. 4D, 5D, 5E and 6C)**. Notably, none of these functional experiments to verify the structural changes were performed in the presence of Fab705, suggesting that these structural changes in our MgtE-Fab complex structure are unlikely to be nonphysiological artifacts due to the use of Fab for structure determination (**Fig. 7**).

Taken together, our structural and functional analyses provide mechanistic insight into the channel-opening mechanisms of MgtE, particularly cytoplasmic pore opening.

## MATERIALS AND METHODS

### Preparation of the MgtE-Fab complex

The full-length *T. thermophilus* MgtE gene was subcloned into a pET28a-derived vector containing N-terminal hexahistidine tags and a human rhinovirus 3C (HRV3C) protease cleavage site, as described previously (Hattori et al. 2007). The protein was overexpressed in *E. coli* C41 (DE3) cells in LB medium containing 30 μg/ml kanamycin at 37 °C by induction at an OD_600_ of ∼0.5 with 0.5 mM isopropyl D-thiogalactoside (IPTG) for 16 hours at 18 °C. The cells were subsequently harvested by centrifugation (6,000 × g, 15 minutes) and resuspended in buffer H [50 mM HEPES (pH 7.0), 150 mM NaCl, 0.5 mM phenylmethanesulfonyl fluoride (PMSF)]. All purification steps were performed at 4 °C. The cells were then disrupted with a microfluidizer, and the supernatants were collected after centrifugation (20,000 × g, 30 minutes) and ultracentrifugation (200,000 × g, 1 hour). Next, the membrane fraction was solubilized with buffer S [50 mM HEPES (pH 7.0), 150 mM NaCl, 2% (w/v) n-dodecyl-beta-d-maltopyranoside (DDM) (Anatrace, USA) 0.5 mM PMSF] for 2 hours. The solubilized fraction was loaded onto a Ni-NTA column pre-equilibrated with buffer A [50 mM HEPES (pH 7.0), 150 mM NaCl, 0.05% (w/v) DDM] containing 20 mM imidazole and mixed for 1 hour. The column was then washed with buffer A containing 50 mM imidazole, and the protein was eluted with buffer A containing 300 mM imidazole. The N-terminal hexahistidine tag was cleaved by HRV3C protease against dialysis buffer (buffer A containing 20 mM imidazole) overnight, and the cleaved protein was reloaded onto a Ni-NTA column pre-equilibrated with dialysis buffer.

The flow-through fraction from the column was collected and concentrated using an Amicon Ultra 50K filter (Merck Millipore) and then loaded onto a Superdex 200 10/300 size-exclusion chromatography (SEC) column (GE Healthcare) in buffer B [20 mM HEPES (pH 7.0), 150 mM NaCl] containing 0.05% (w/v) DDM. The main peak fractions from the column were collected and mixed with PMAL-C8 amphipol (Anatrace) at a ratio of 1:5 (w:w) overnight. The sample was mixed with Bio-Beads SM-2 (Bio-Rad) and rotated gently for 4 hours. The Bio-Beads were subsequently removed by a disposable Poly-Prep column, and the eluents were concentrated using an Amicon Ultra 50K filter and loaded onto a Superdex 200 10/300 SEC column in buffer B. The SEC fractions corresponding to the amphipol-reconstituted MgtE were collected, mixed with the Fab antibody fragment Fab705 at a ratio of 1:2 (w:w) for 1 hour, and applied to a Superdex 200 10/300 SEC column in buffer B. The SEC fractions corresponding to the MgtE-Fab complex were collected and concentrated to ∼1 mg/ml with an Amicon Ultra 50K filter for single-particle analysis using cryo-EM.

### EM data acquisition and analysis

A total of 4.0 μl of the MgtE-Fab complex in PMAL-C8 was applied to a glow-discharged holey carbon-film grid (Zhongkexinghua BioTech, GiG-A31213, 300M-Au-R1.2/1.3) blotted with a Vitrorobot (FEI) system using a 1.0-s blotting time with 100% humidity at 9 °C and plunge-frozen in liquid ethane. Cryo-EM images were collected on a Titan Krios (FEI) electron microscope operated at an acceleration voltage of 300 kV. The specimen stage temperature was maintained at 80 K. Images were recorded with a K2 Summit direct electron detector camera (Gatan Inc.) set to superresolution mode with a pixel size of 0.41 Å (a physical pixel size of 0.82 Å) and a defocus ranging from −1.5 µm to −2.3 µm. The dose rate was 8 e^-^ s^−1^, and each movie was 6 s long, dose-fractioned into 40 frames with an exposure of 1.75 e^-^ Å^−2^ for each frame. The cryo-EM data are summarized in **Table S2**.

### Image processing

A total of 5,610 movies of the MgtE-Fab complex were motion-corrected and binned with MotionCor2 (Zheng et al. 2017) with 5 × 5 patches, producing summed and dose-weighted micrographs with a pixel size of 0.82 Å. Contrast transfer function (CTF) parameters were estimated by CTFFIND 4.1 (Rohou and Grigorieff 2015). Particle picking and further image processing were performed using RELION 3.0 (Zivanov et al. 2018). A total of 1,860,461 particles were autopicked and extracted with a box size of 256 × 256 pixels. After several rounds of 2D classification, the re-extracted particles were classified into 12 classes for 3D classification without any symmetry imposed. Refinement and postprocessing using RELION produced a final map at a 3.7 Å resolution from 168,911 particles with C2 symmetry imposed. The final 3D reconstruction was calculated from 2 × 2 binned images (0.82 Å). The final resolution was estimated using the Fourier shell correlation (FSC) = 0.143 criterion on the corrected FSC curves, in which the influence of the mask was removed. The local resolution was estimated using RELION. The workflow for image processing and for the 3D reconstruction and angular distribution plot are shown in **Fig. S3**.

### Model building

The initial model of Mg^2+^-free MgtE-Fab was manually built starting from the MgtE TM domain structure and the homology model of Fab705 generated by SWISS-MODEL (Biasini et al. 2014). Manual model building was performed using COOT (Emsley et al. 2010). Real-space refinement was performed using PHENIX software (Afonine et al. 2012). The final atomic model of MgtE includes residues 271-448 for chains A and B. All figures showing structures were generated using PyMol (https://pymol.org/). The sequence alignment figure was generated using Clustal Omega (Sievers and Higgins 2017) and ESPript 3.0 (Robert and Gouet 2014). The MgtE pore shown in **Fig. 4** was calculated by HOLE (Smart et al. 1996).

### Preparation of antibody fragments

Mouse monoclonal antibodies against MgtE were raised by an established method (Jaenecke et al. 2018). Briefly, a proteoliposome antigen was prepared by reconstituting purified, functional MgtE at high density in phospholipid vesicles consisting of a 10:1 mixture of egg phosphatidylcholine (Avanti Polar Lipids) and the adjuvant lipid A (Sigma) to facilitate the immune response. BALB/c mice immunization was performed with the proteoliposome antigen using three injections at two-week intervals.

Antibody-producing hybridoma cell lines were generated using a conventional fusion protocol (KÖHLER and MILSTEIN 1975). Hybridoma clones producing antibodies recognizing conformational epitopes in MgtE were selected by an enzyme-linked immunosorbent assay on immobilized phospholipid vesicles containing purified MgtE (liposome ELISA), allowing positive selection of the antibodies that recognized the native conformation of MgtE. Additional screening for reduced antibody binding to sodium dodecyl sulfate (SDS)-denatured MgtE was used for negative selection against linear epitope-recognizing antibodies. Stable complex formation between MgtE and each antibody clone was checked using fluorescence-detection SEC (FSEC). From these screens, we isolated four MgtE-specific monoclonal antibodies: YN0705, YN0710, YN0721 and YN0736. Whole IgG molecules, collected from the large-scale culture supernatant of monoclonal hybridomas and purified using protein G affinity chromatography, were digested with papain, and Fab fragments were isolated using a HiLoad 16/600 Superdex 200 gel filtration column followed by protein A affinity chromatography. The cDNAs encoding the light and heavy chains of Fab YN0705 were cloned from hybridoma cells using rapid amplification of 5’ complementary DNA ends (5’-RACE) and sequenced.

### Isothermal titration calorimetry

The binding affinities of the Fab antibody fragment Fab705 for full-length MgtE were measured using MicroCal iTC200 (GE Healthcare). The full-length MgtE proteins were purified by a similar procedure to that described above but with SEC buffer [buffer B containing 0.05% (w/v) DDM] containing either 0 or 20 mM MgCl_2_. Fab was also used for gel filtration with the same SEC buffer containing either 0 or 20 mM MgCl_2_. The peak fractions of MgtE were collected and diluted to approximately 2.5 µM (monomer) with SEC buffer, while Fab was diluted to 25 µM. Fab was injected into the cell containing the full-length MgtE 20 times (0.5 µl for injection 1, 2 µl for injections 2-20) at 25 °C, with 120-s intervals between injections. The background data were measured by injecting the SEC buffer into a cell containing the full-length MgtE. The results were analyzed by Origin8 software (MicroCal). The experiments were repeated three times for each measurement, and similar results were achieved.

### Chemical cross-linking

The wild-type and mutant MgtE proteins were purified as described above, except that dithiothreitol (DTT) was added at a final concentration of 20 mM to the MgtE mutant proteins before removal by size exclusion chromatography, and then diluted to 0.5 mg/ml with buffer B containing 0.05% (w/v) DDM. MgtE proteins (4.0 μl) were mixed with 0.5 µl of MgCl_2_ at appropriate concentrations and incubated for 30 minutes at 4 °C. Then, 0.5 µl of 10 mM Cu^2+^ bis-1,10-phenanthroline was added at a 1:3 molar ratio, and the samples were incubated for another 30 minutes at 4 °C. The reaction mixtures were loaded with nonreduced loading buffer and analyzed immediately by sodium dodecyl sulfate-polyacrylamide gel electrophoresis (SDS-PAGE). The relative intensity of the bands was analyzed by ImageJ software and fitted to a sigmoidal equation. The experiments were repeated six times.

### Molecular dynamics simulations

As described previously (Wang et al. 2018)(Li et al. 2018), DESMOND (Shaw 2005) was used to perform MD analysis with a constant number of particles, constant pressure (1 bar) and temperature (300 K) (NPT) and periodic boundary conditions (PBCs) by using the Nose-Hoover chain thermostat and Martyna-Tobias-Klein barostat methods. The MgtE TM domain dimer was embedded in a phosphoryl-oleoyl phosphatidylcholine (POPC) bilayer. The structure of MgtE/POPC was solvated in an orthorhombic box of SPC water molecules. All-atom OPLS_2005 force fields for proteins, ions, lipids and simple point charge (SPC) waters were used in all simulations (Kaminski et al. 2001)(Jorgensen et al. 1996). Before performing simulations, a default relaxation protocol in DESMOND was employed: (1) NVT ensemble with Brownian dynamics for 100 ps at 10 K with small-time step, and solute heavy atom restrained; (2) NVT ensemble using Berendsen thermostat for 12 ps at 10 K with small-time step, and solute heavy atom restrained; (3) NPT ensemble using a Berendsen thermostat and barostat for 12 ps at 10 K and 1 atm with solute heavy atom restrained; (4) NPT ensemble using a Berendsen for 12 ps at 300 K and 1 atm with solute heavy atom restrained; (5) NPT ensemble using a Berendsen for 24 ps at 300 K and 1 atm with no restraints. After the relaxation, MD simulation was performed for 200 ns. The integration time step used was 2 fs, and coordinate trajectories were saved every 200 ps. A DELL T7910 graphics work station (with multiple NVIDA Tesla K40C GPUs) was used to run the MD simulations, and a 12-CPU CORE DELL T7500 graphics work station was used to perform the preparation, analysis and visualization. The Simulation Event Analysis module in DESMOND was used to analyze MD trajectories.

### Electrophysiological recordings

Patch clamp recording of MgtE using *E. coli* giant spheroplasts was performed in the inside-out configuration, as described previously (Martinac et al. 1987)(Hattori et al. 2009). In brief, spheroplasts were placed in a bath solution containing 210 mM N‐methyl‐D‐glucamine, 90 mM MgCl_2_, 300 mM D-glucose, and 5 mM HEPES (pH 7.2). After gigaseal formation with a borosilicate glass pipette (Harvard Apparatus, Kent, UK) with 6-8 mOhm resistance, the bath medium was changed to bath solution containing 300 mM N‐methyl‐D‐glucamine, 300 mM D-glucose, and 5 mM HEPES (pH 7.2) by perfusion using a Rainer perfusion pump and a custom-made perfusion system. Bath medium containing 2 μM Fab705 was added by perfusion. The pipette solution contained 210 mM N‐methyl‐D‐glucamine, 90 mM MgCl_2_, 300 mM sucrose, and 5 mM HEPES (pH 7.2). Currents were measured with an Axopatch 200B amplifier and recorded with a Digidata 1440B A/D converter under the control of pCLAMP software (Molecular Devices). Currents were filtered at 2 kHz and recorded at 5 kHz.

### *E. coli* genetics

As described previously (Hattori et al. 2009), the Mg^2+^-auxotrophic *E. coli* strain (BW25113 Δ*mgtA* Δ*corA* Δ*yhiD* DE3) was transformed with MgtE or MgtE mutant plasmids and grown on LB medium plates containing 50 μg/ml kanamycin and 100 mM MgSO_4_. Each transformant was inoculated in liquid LB medium containing 50 μg/ml kanamycin and 100 mM MgSO_4_ and incubated at 37 °C overnight with shaking. The liquid samples were diluted 1:100 with the same medium and grown until the OD600 reached 0.6. The sample cultures were then serially diluted 10-fold in LB medium containing 50 μg/ml kanamycin, spotted onto assay plates and incubated at 37 °C overnight, as indicated in **Fig. 6C**.

In the Ni^2+^ sensitivity assay, the isogenic magnesium-prototrophic wild-type *E. coli* strain (W3110 DE3) was transformed with the plasmid, and transformants were obtained on LB (+50 μg/ml kanamycin) plates. The liquid samples were diluted 1:100 with the same medium and grown until the OD600 reached 0.6. The sample cultures were then serially diluted 10-fold and spotted onto assay plates, and the plates were incubated at 37 °C overnight, as indicated in **Fig. 4D**.

The expression of MgtE and its mutants was confirmed by Western blotting (**Fig. S8 and S9)**.

### Western blot analysis

The Mg^2+^-auxotrophic *E. coli* strain (BW25113 Δ*mgtA* Δ*corA* Δ*yhiD* DE3) was transformed with the plasmids, and transformants were obtained on LB (+50 μg/ml kanamycin) plates supplemented with 100 mM MgSO_4_. Each transformant was cultured in LB liquid medium supplemented with 50 μg/ml kanamycin and 100 mM MgSO_4_ overnight. The *E. coli* cell numbers were adjusted based on the OD600. Whole-cell extracts were prepared, resolved by SDS-PAGE (10% polyacrylamide), and transferred to Hybond-ECL (GE Healthcare). The transferred proteins were stained with an anti-His6 polyclonal antibody, anti-His-tag-HRP direct T (MBL). Antibody-bound proteins were visualized using Western blot detection reagents (Immunostar LD, Wako Laboratory Chemicals). Images were captured with an LAS-3000 Mini imaging system (Fujifilm).

### FSEC experiments

Purified full-length MgtE and Fab705 proteins were diluted to 1 mg/ml with buffer B containing 0.05% (w/v) DDM. The MgtE-Fab complex was formed by mixing MgtE and Fab at mass ratios of 1:0, 0:1, 1:0.5, 1:1 and 1:2 in the presence of 0 mM or 20 mM Mg^2+^. The samples were then loaded onto a Superdex 200 Increase 10/300 size-exclusion chromatography (SEC) column (GE Healthcare, USA) connected to an RF-20Axs fluorescence detector (Shimadzu, Japan) (excitation: 280 nm, emission: 325 nm) with buffer B containing 0.05% (w/v) DDM and 0 mM or 20 mM Mg^2+^.

The complex disruption experiment was performed by adding Mg^2+^ at a final concentration of 20 mM to the preformed MgtE-Fab complex, which was prepared by mixing MgtE and Fab705 at a mass ratio of 1:2 in 0 mM Mg^2+^ and loading the sample on an FSEC column with buffer B containing 0.05% (w/v) DDM and 20 mM Mg^2+^.

### Data availability

The atomic coordinates and structural factors of the MgtE-Fab structure were deposited in the Protein Data Bank. The accession numbers for the MgtE-Fab structure are PDB: 6LBH and EMD: EMDB-0869. All other data are available from the authors upon reasonable request.

## ACKNOWLEDGMENTS

We thank the staff scientists at the Center for Biological Imaging, Institute of Biophysics, and National Center for Protein Science Shanghai (Chinese Academy of Sciences) for technical support with cryo-EM data collection (project numbers: CBIapp20180107; CBIapp201807006; CBIapp201807007; 2017-NFPS-PT-001632; 2018-NFPS-PT-002187), Yumi Sato for technical assistance in the generation of antibodies, Namba laboratory members for technical support with cryo-EM data collection, Dr Hideaki E. Kato (University of Tokyo) for critical comments on the manuscript and Drs. Chia-Hsueh Lee (St. Jude Children’s Research Hospital) and Muneyoshi Ichikawa (NAIST) for their technical advice on cryo-EM data processing. This work was supported by funding from the Ministry of Science and Technology of China (2016YFA0502800) to M.H.; funding from the National Natural Science Foundation of China (projects 31570838 and 31850410466) to M.H., the Innovative Research Team of High-level Local Universities in Shanghai, and Shanghai Key Laboratory of Bioactive Small Molecules (ZDSYS14005); grants from Basis for Supporting Innovative Drug Discovery and Life Science Research (BINDS) from the Japan Agency of Medical Research and Development (AMED) (grant no. 19am0101079; Support No. 0451); Research on Development of New Drugs from the AMED; and Grants-in-Aid for Scientific Research from the Japan Society for the Promotion of Science (JSPS) (nos. 18K05334 and 19H00923).

## ABBREVIATIONS

cryo-EM: cryoelectron microscopy
MgtE: Magnesium transporter E
SLC: Solute carrier
CBS: cystathionine beta synthase
TM: transmembrane
Fab: fragment antigen-binding
FSEC: fluorescence-detection size-exclusion chromatography
ITC: isothermal titration calorimetry
DDM: n-dodecyl-beta-d-maltopyranoside
HS-AFM: high-speed atomic force microscopy

## AUTHOR CONTRIBUTIONS

F.J. expressed and purified MgtE with assistance from M.S. and performed cryo-EM experiments with assistance from S.S., and J.M., T.F., Y.M., and K.N. contributed to the early stage of cryo-EM experiments. F.J. and M.H. performed model building. F.J. conducted ITC and chemical cross-linking experiments. Y.N., K.L., T.U., Y.N., N.N. and S.I. generated the anti-MgtE-Fab antibody. H.T., A.T., T.N., R.I. and O.N. contributed to the initial trial of MgtE crystallization under Mg^2+^-free conditions. T.K. prepared giant spheroplasts of *E. coli* for patch clamp recording. A.M. performed the electrophysiology work. M.W. and K.I. performed the *E. coli* genetics work. J.W. and Y. Y performed the MD simulation. F.J. and M.H. wrote the manuscript. M.H. supervised the research. All authors discussed the manuscript.

## COMPLIANCE WITH ETHICS GUIDELINES

Fei Jin, Minxuan Sun, Takashi Fujii, Yurika Yamada, Jin Wang, Andrés D. Maturana, Miki Wada, Shichen Su, Jinbiao Ma, Hironori Takeda, Tsukasa Kusakizako, Atsuhiro Tomita, Yoshiko Nakada-Nakura, Kehong Liu, Tomoko Uemura, Yayoi Nomura, Norimichi Nomura, Koichi Ito, Osamu Nureki, Keiichi Namba, So Iwata, Ye Yu and Motoyuki Hattori declare that they have no conflict of interest. All institutional and national guidelines for the care and use of laboratory animals were followed.

## Notes

### Competing Interest Statement

The authors have declared no competing interest.

