## supplemental figures for "Cryo-EM structure of the MgtE Mg^2+^ channel pore domain in Mg^2+^-free conditions reveals cytoplasmic pore opening"

### Supplementary Materials

#### Supplementary Figure legends

**Figure S1 FSEC analysis of MgtE-Fab complex formation.**

(**A, B**) FSEC analysis of MgtE-Fab complex formation in the absence of $MgCl_2$ (**A**) and presence of 20 mM $MgCl_2$. (**B**). The MgtE-Fab complex was formed by mixing MgtE and Fab at the indicated mass ratios. (**C**) The complex disruption experiment was performed by adding $Mg^{2+}$ at a final concentration of 20 mM to the preformed MgtE-Fab complex, which was prepared by mixing MgtE and Fab705 at a mass ratio of 1:2 in 0 mM $Mg^{2+}$. FSEC analysis was performed in the presence of 20 mM $MgCl_2$ (**C**).

**Figure S2 Size-exclusion chromatography and SDS-PAGE of the Fab-MgtE complex**

(**A**) Size-exclusion chromatography of the MgtE-Fab complex. The former, middle and latter peaks were void, Fab-MgtE complex and free Fab, respectively. The fractions from the 10.5 to 12.0 ml elution positions were pooled as a cryo-EM sample. (**B**) SDS-PAGE of the SEC fractions and the cryo-EM sample.

**Figure S3 Flowchart for cryo-EM data processing**

(A) Overview of the data processing workflow, including particle picking, classification, and 3D refinement. All processing steps were performed in RELION. (B) Euler angle distribution plot of all particles included in the calculation of the MgtE-Fab complex, with C2 symmetry imposed.

**Figure S4 EM density map.**

(**A-E**) Representative EM density maps contoured at 4.0 σ. The TM1 helix (residues 278-310) (A), TM2 helix (residues 315-340) (B), TM3 helix (residues 353-380) (C), TM4 helix (residues 384-416) (D), and TM5 helix (residues 421-448) (E) in chain A are shown as stick representations.

**Figure S5 Fab-binding site.**

A close-up view of the MgtE-Fab interface on the cytoplasmic side (**A**) and of the corresponding region in full-length MgtE in the $Mg^{2+}$-bound form (**B**).

**Figure S6 Structural comparisons.**

Superimpositions of our cryo-EM structure of the MgtE-Fab complex under $Mg^{2+}$-free conditions and $Mg^{2+}$-free cytoplasmic domain structure onto full-length MgtE in the $Mg^{2+}$-bound form, viewed parallel to the membrane (A) and from the cytoplasmic side (**B**). The coloring scheme of the full-length MgtE structure is the same as that in Figure 1. The $Mg^{2+}$-free cytoplasmic domain structure is colored gray. The TM domain and Fabs in the MgtE-Fab complex are colored red and orange, respectively.

**Figure S7 MD simulations.**

(**A**) Structural deviation from the $Mg^{2+}$-free MgtE TM domain structure during the 2-μs MD simulation. (**B-D**) Cα distances between Thr336 (chain B) and Leu421 (chain A), between Thr336 (chain A) and Leu421 (chain B), and between Ala317 (chains A and B) during the 2-μs MD simulation.

**Figure S8 Western blot of the MgtE N424A mutant.**

**Figure S9 Western blot of the MgtE glycine mutants.**

**Video S1. Structural changes in the MgtE TM domain.**

**Table S1. ITC statistics**

**Table S2. Cryo-EM data collection, refinement and validation statistics**

A

B

C

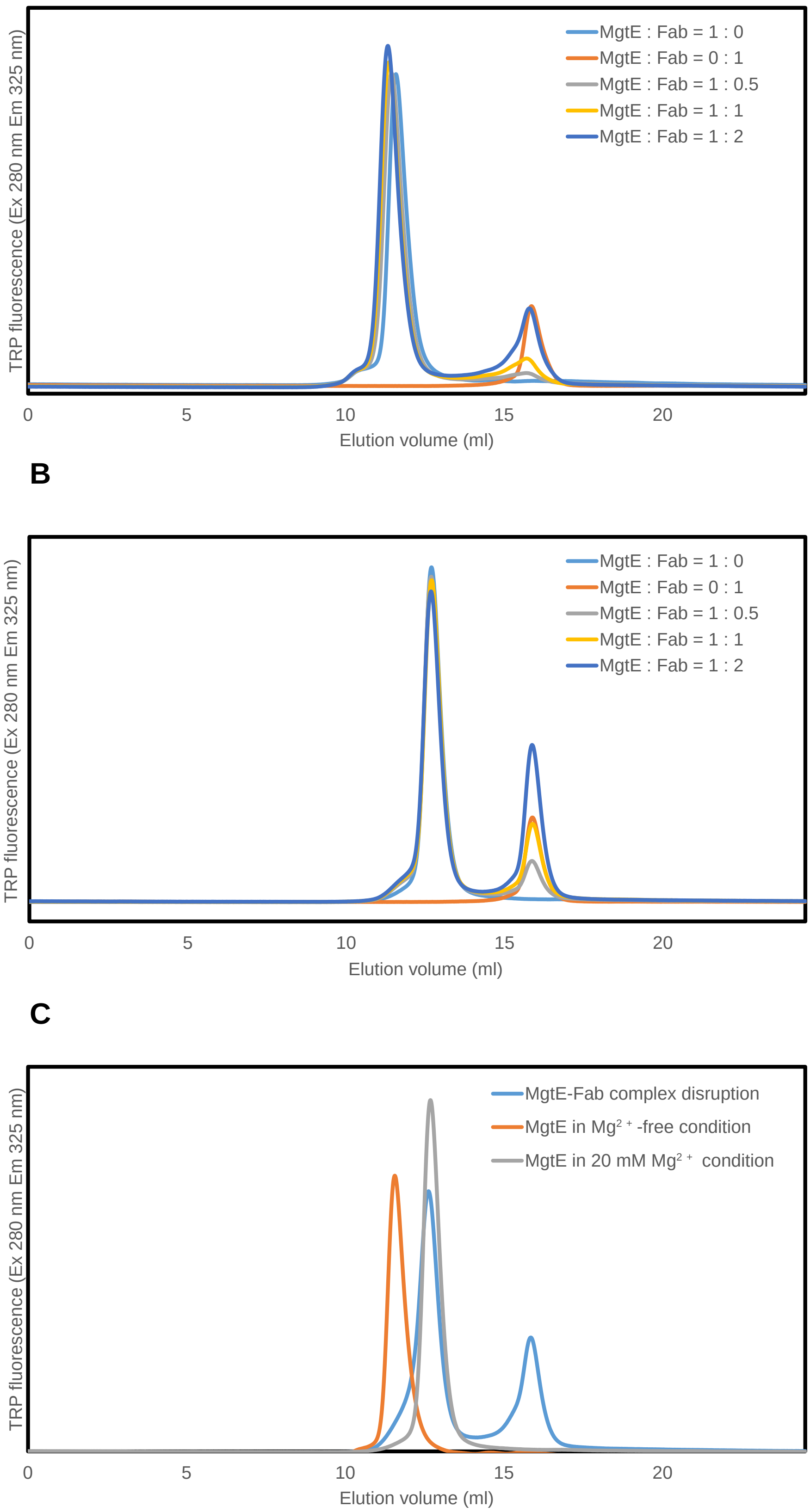

### Figure S1

A

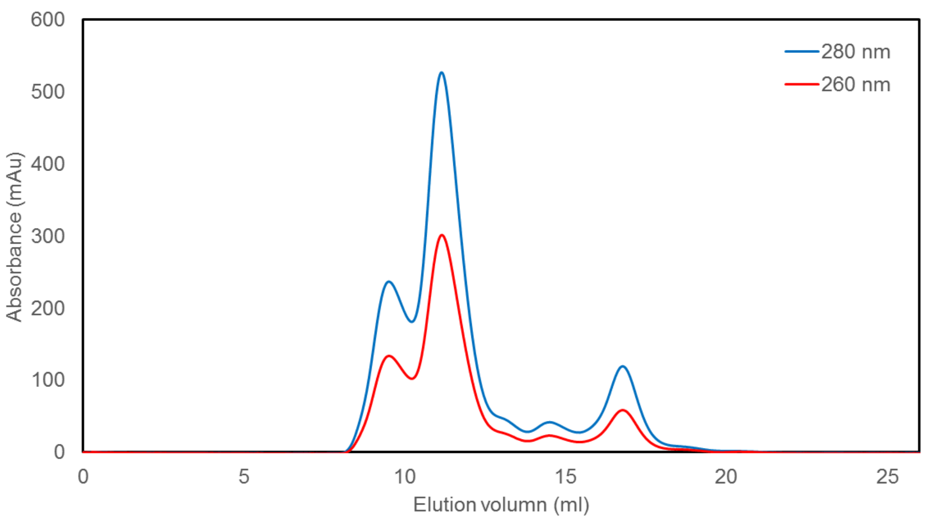

600
500
400
300
200
100
0
Absorbance (mAu)
0
5
10
15
20
25
Elution volumn (ml)
280 nm
260 nm

B

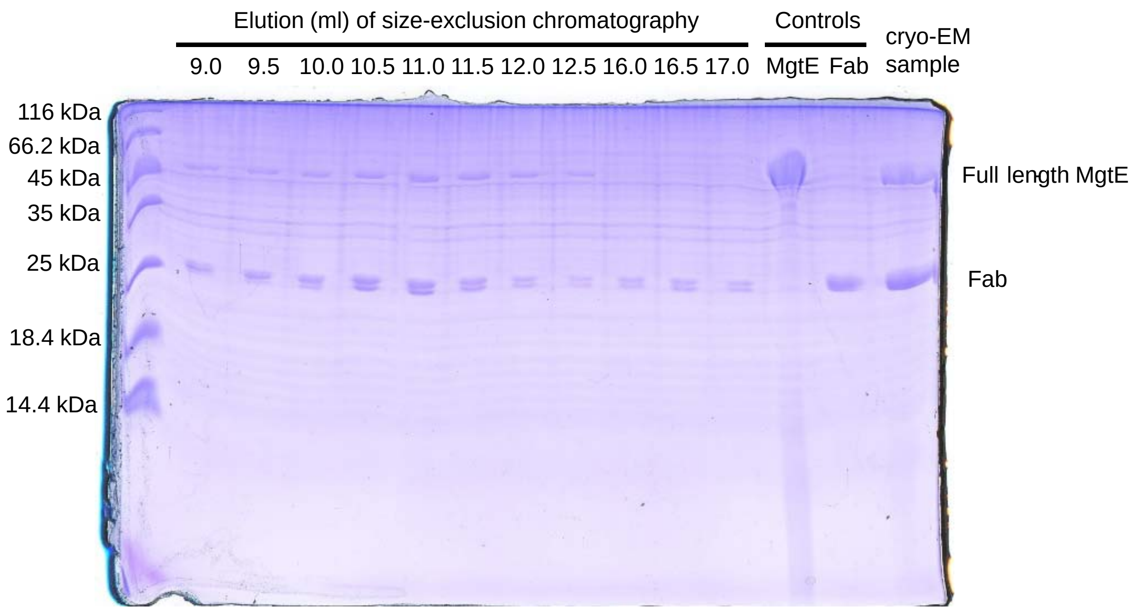

Elution (ml) of size-exclusion chromatography
Controls
cryo-EM sample
9.0 9.5 10.0 10.5 11.0 11.5 12.0 12.5 16.0 16.5 17.0
MgtE Fab
116 kDa
66.2 kDa
45 kDa
35 kDa
25 kDa
18.4 kDa
14.4 kDa
Full length MgtE
Fab

Figure S2

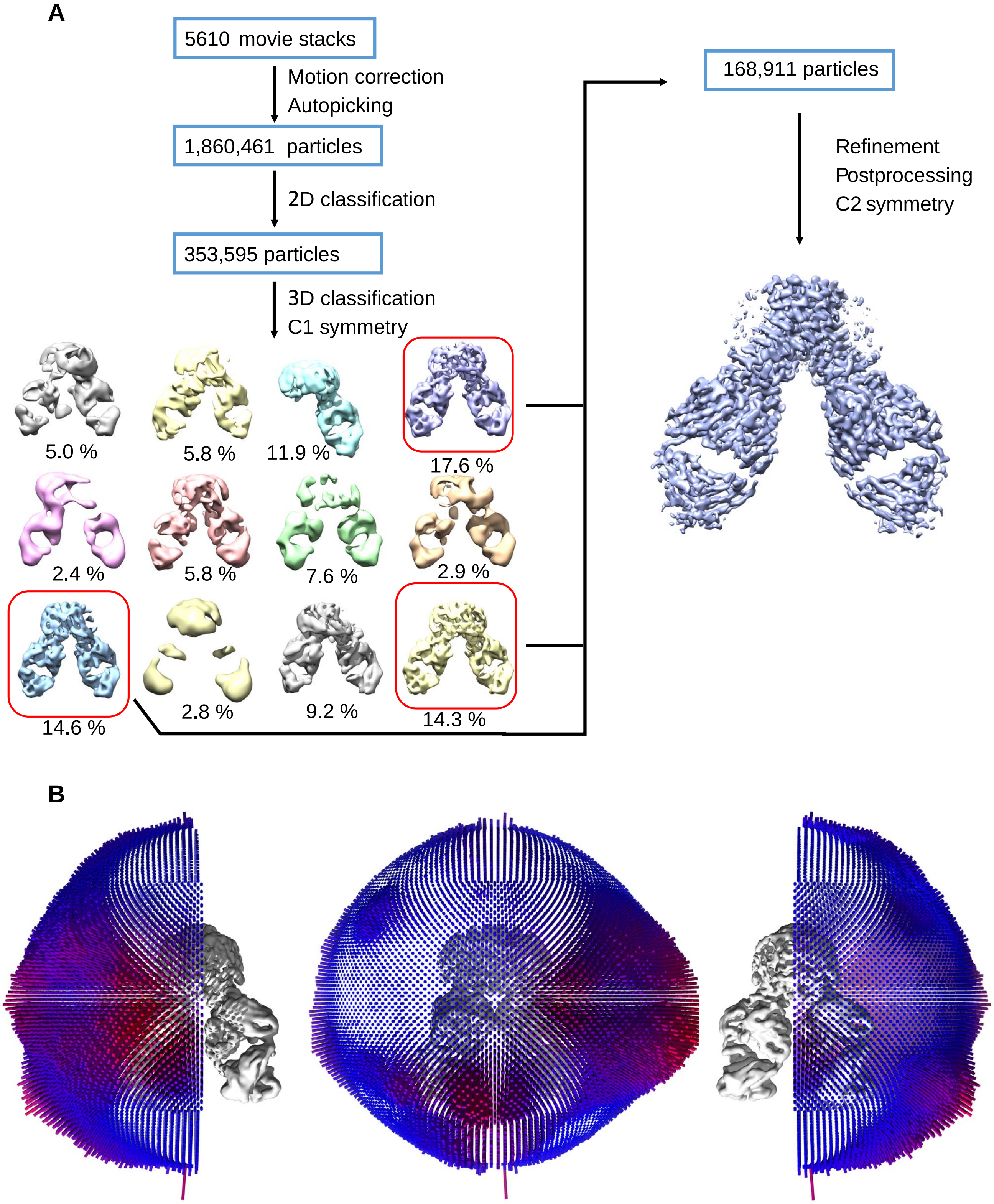
A
5610 movie stacks
Motion correction
Autopicking
1,860,461 particles
2D classification
353,595 particles
3D classification
C1 symmetry
5.0 %
5.8 %
11.9 %
17.6 %
2.4 %
5.8 %
7.6 %
2.9 %
14.6 %
2.8 %
9.2 %
14.3 %
168,911 particles
Refinement
Postprocessing
C2 symmetry
B

Figure S3

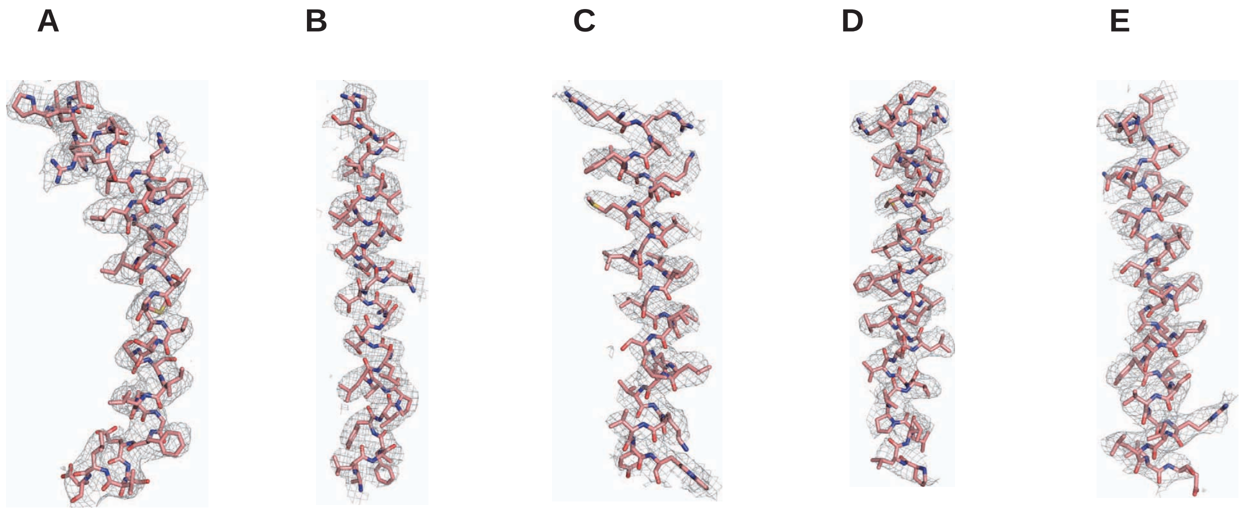

Figure S4

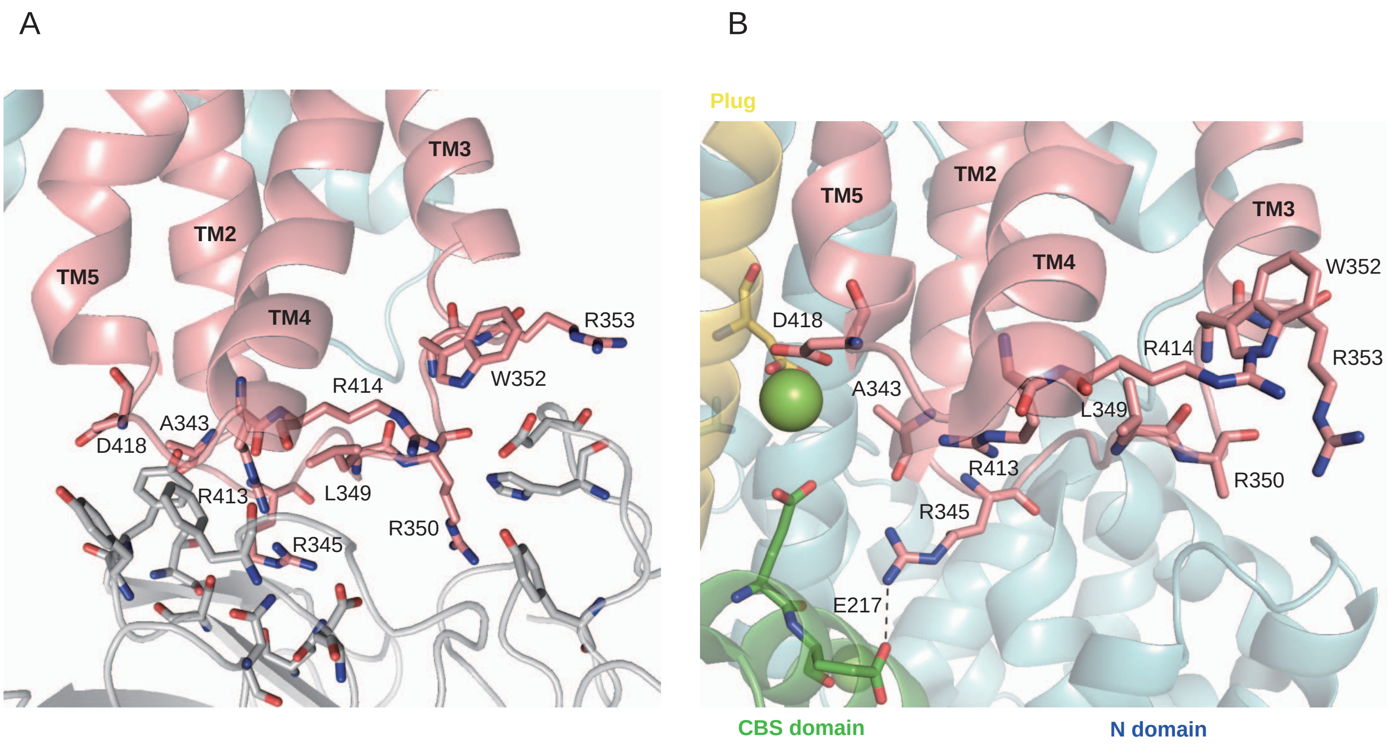
A
TM3
TM2
TM5
TM4
R353
R414
W352
A343
D418
R413
L349
R345
R350
B
Plug
TM5
TM2
TM3
TM4
W352
D418
R414
R353
A343
L349
R413
R350
R345
E217
CBS domain
N domain

Figure S5

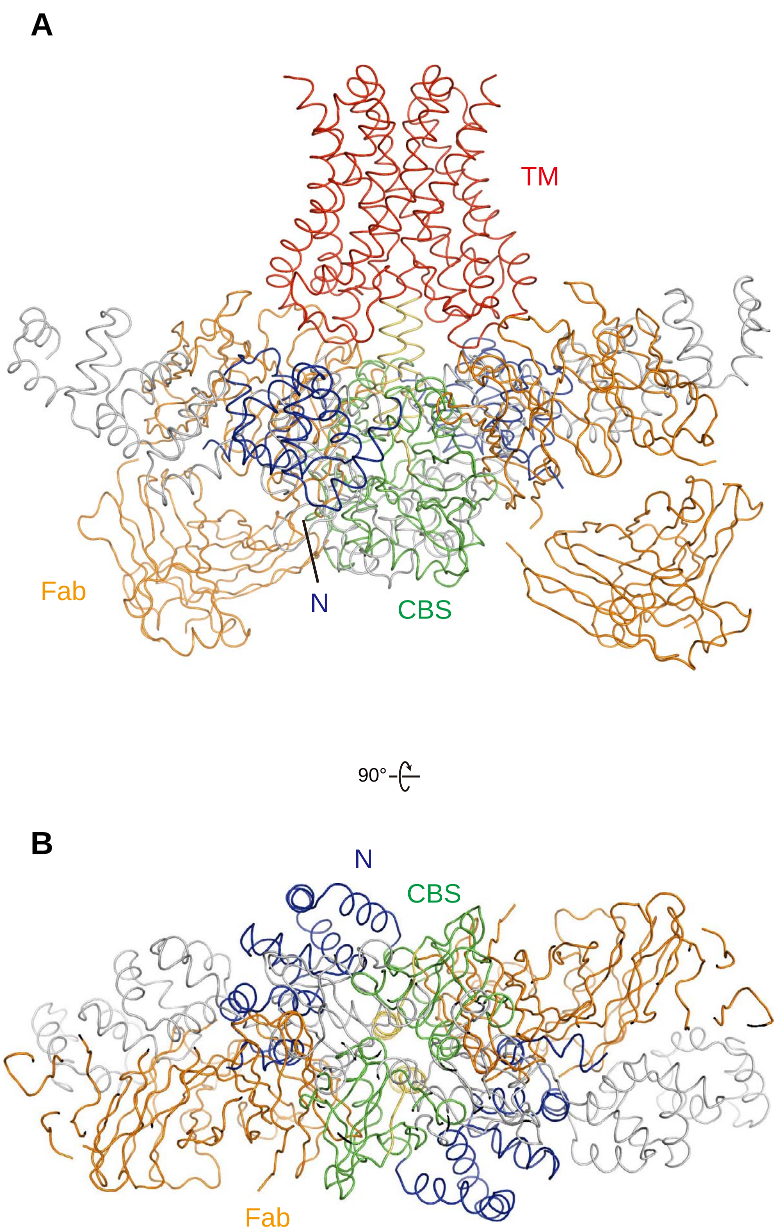

Figure S6

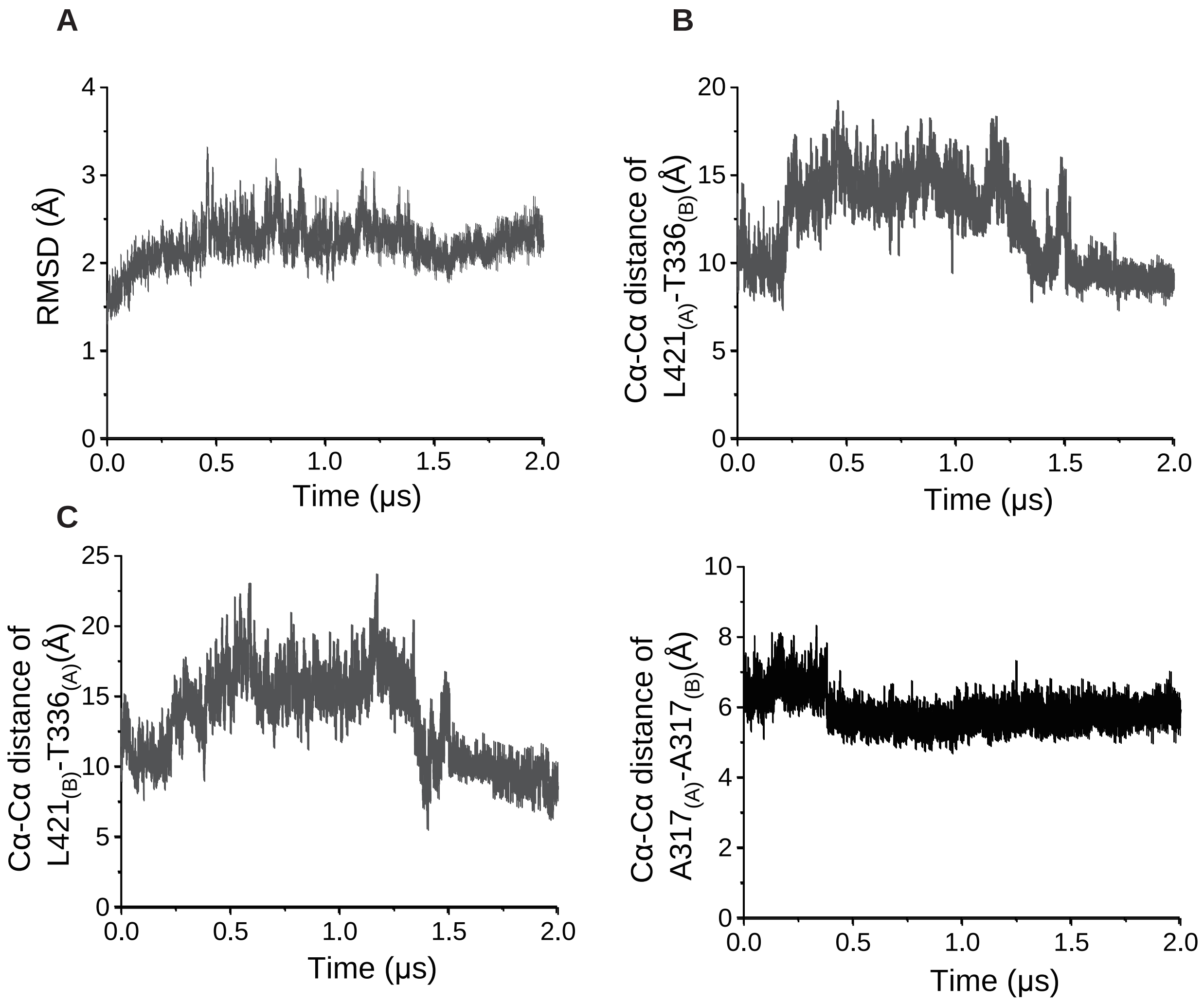
A
B
C
RMSD (Å)
Time (μs)
Cα-Cα distance of L421(A)-T336(B)(Å)
Cα-Cα distance of L421(B)-T336(A)(Å)
Cα-Cα distance of A317(A)-A317(B)(Å)

Figure S7

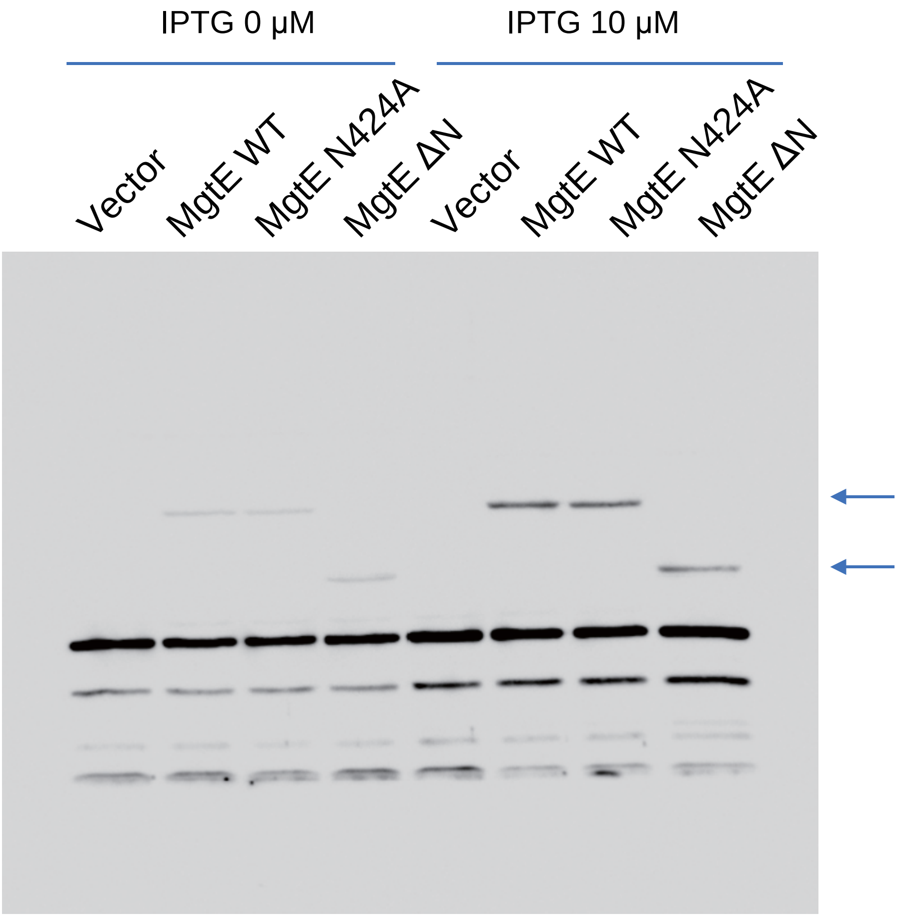
IPTG 0 µM
IPTG 10 µM
Vector
MgtE WT
MgtE N424A
MgtE ΔN
Vector
MgtE WT
MgtE N424A
MgtE ΔN

Figure S8

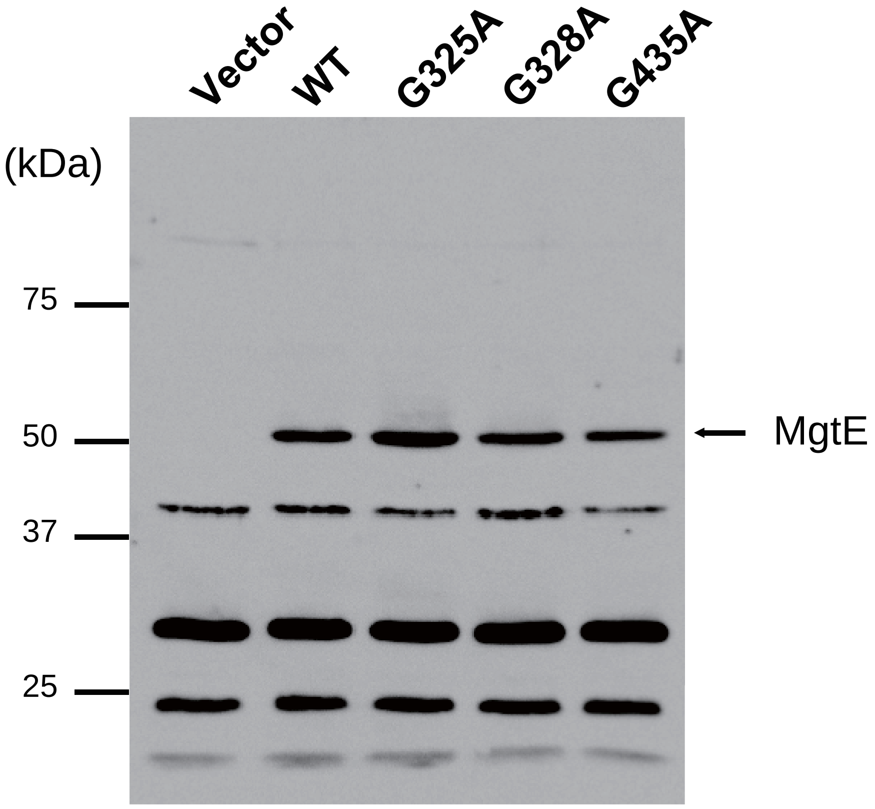

Figure S9

**Table S1. ITC statistics**

| | N | ΔH° (kcal/mol) | −TΔS° (kcal/mol) | ΔG° (kcal/mol) | $K_D$ (μM) |
|---|---|---|---|---|---|
| Without $Mg^{2+}$ | 1.32 ± 0.03 | -9.16 ± 0.31 | 0.11 | -9.05 ± 0.31 | 252.6 ± 71.1 |
| With $Mg^{2+}$ | | | | | ND |

**Table S2. Cryo-EM data collection, refinement and validation statistics**

| | MgtE-Fab (EMDB-0869) (PDB 6LBH) |
|---|---|
| **Data collection and processing** | |
| Magnification | 29,000x |
| Voltage (kV) | 300 |
| Electron exposure (e–/Å$^2$) | 70 |
| Defocus range (μm) | -1.5 to -2.3 |
| Pixel size (Å) | 0.82 |
| Symmetry imposed | *C*2 |
| Initial particle images (no.) | 1,860,461 |
| Final particle images (no.) | 168,911 |
| Map resolution (Å) | 3.7 |
| FSC threshold | 0.143 |
| Map resolution range (Å) | 2.84-4.5 |
| **Refinement** | |
| Initial model used (PDB code) | This study |
| Model resolution (Å) | 3.7 |
| FSC threshold | 0.143 |
| Model resolution range (Å) | 2.84-4.5 |
| Map sharpening *B* factor (Å$^2$) | -167 |
| Model composition | |
| Non-hydrogen atoms | 9,234 |
| Protein residues | 1,236 |
| Ligands | NA |
| *B* factors (Å$^2$) | |
| Protein | 55.87 |
| Ligand | NA |
| R.m.s. deviations | |
| Bond lengths (Å) | 0.010 |
| Bond angles (°) | 1.069 |
| Validation | |
| MolProbity score | 2.26 |
| Clashscore | 9.97 |
| Poor rotamers (%) | 2.37 |
| Ramachandran plot | |
| Favored (%) | 92.9 |
| Allowed (%) | 6.9 |
| Disallowed (%) | 0.2 |
